# Rescue of hippocampal synaptic plasticity and memory performance by Fingolimod (FTY720) in APP/PS1 model of Alzheimer’s disease is accompanied by correction in metabolism of sphingolipids, polyamines, and phospholipid saturation composition

**DOI:** 10.1101/2025.01.17.633452

**Authors:** Karel Kalecký, Luna Buitrago, Juan Marcos Alarcon, Abanish Singh, Teodoro Bottiglieri, Rima Kaddurah-Daouk, Alejandro Iván Hernández, Alzheimer’s Disease Metabolomics Consortium

## Abstract

Previously, our metabolomic, transcriptomic, and genomic studies characterized the ceramide/sphingomyelin pathway as a therapeutic target in Alzheimer’s disease, and we demonstrated that FTY720, a sphingosine-1-phospahate receptor modulator approved for treatment of multiple sclerosis, recovers synaptic plasticity and memory in APP/PS1 mice. To further investigate how FTY720 rescues the pathology, we performed metabolomic analysis in brain, plasma, and liver of trained APP/PS1 and wild-type mice. APP/PS1 mice showed area-specific brain disturbances in polyamines, phospholipids, and sphingolipids. Most changes were completely or partially normalized in FTY720-treated subjects, indicating rebalancing the “sphingolipid rheostat”, reactivating phosphatidylethanolamine synthesis via mitochondrial phosphatidylserine decarboxylase pathway, and normalizing polyamine levels that support mitochondrial activity. Synaptic plasticity and memory were rescued, with spermidine synthesis in temporal cortex best corresponding to hippocampal CA3-CA1 plasticity normalization. FTY720 effects, also reflected in other pathways, are consistent with promotion of mitochondrial function, synaptic plasticity, and anti-inflammatory environment, while reducing pro-apoptotic and pro-inflammatory signals.

## Introduction

Alzheimer’s disease (AD) is the most common neurodegenerative disease, which, along with related dementias, affects over 50 million people globally^1^. From a neuropathology perspective, AD has two major characteristics: (i) accumulations of extracellular aggregated peptide known as beta amyloid (Aβ), which form the well-characterized senile plaques; and (ii) intracellular accumulation of an abnormally phosphorylated protein, tau, leading to the formation of neurofibrillary tangles^2–4^. AD is characterized by a progressive decline of function in multiple cognitive domains, including episodic memory, speech, visuospatial processing, and executive function, that eventually impairs independence and affects activities of daily living that, in the late stage of the disease, require caregiver assistance^5^. The cost of care for individuals with AD is estimated at 1 trillion USD annually, a figure that is expected to double in the next decade^5^. Thus, the search for the discovery of biomarkers to help with early diagnosis and effective therapies have become increasingly relevant to address the expansion of the disease and the economic problem.

Although AD is a multifactorial disease^6^ associated with changes in multiple metabolic pathways^7,8^, growing evidence implicates dysregulation of lipid metabolism in AD pathogenesis^9–15^. More than half of the genes identified in genome-wide association studies are linked to lipid homeostasis^10^. This includes apolipoprotein E allele ε4, one of the strongest risk factors for developing late onset sporadic AD^16^. A recent study combining positron emission tomography (PET) for tau and Aβ with human gene expression data showed several lipid metabolism-related genes to be involved in the spread of tau and Aβ in AD patients^17^.

Lipids comprise 50-60% of the brain’s dry weight and are the basic structural component of neuronal cell membranes^18^. Changes in lipid homeostasis in the nervous system are associated with cell death, loss of synaptic plasticity, and, ultimately, neurodegeneration^19–21^. Sphingolipids, a complex class of lipids highly enriched in the central nervous system (CNS), are composed of a long-chain sphingoid base backbone (sphingosine), linked to a fatty acid via an amide bond, and one of various types of polar head groups. Sphingolipids are associated with signal transmission, membrane structure, and myelin stability. Some members of the sphingolipid family, such as ceramide, sphingosine, sphingosine-1-phosphate (S1P), and ceramide-1-phosphate (C1P), are bioactive signaling molecules playing a major role in cell growth, differentiation, apoptosis, senescence, survival, migration, and inflammation^22^. While ceramide and sphingosine elevation induces apoptosis, S1P is required for cell proliferation and survival. The balance between S1P and ceramide/sphingosine is referred to as the “sphingolipid rheostat” and appears to be the central mechanism for determining cellular fate^23,24^.

Recent discoveries suggest that subtle changes in the sphingolipid balance could be involved in AD. Several studies report that ceramides are elevated in brain tissue of AD patients compared to age-matched individuals^25–28^, and similarly for sphingosine^29^, while sphingomyelin and S1P are decreased^28,30^. Furthermore, lipidomic studies revealed increased ceramide and decreased sphingomyelin levels in plasma of AD patients compared to cognitively normal controls^31^. Longitudinal studies in AD patients reported that distinct sphingolipid species were consistently associated with brain atrophy, cognitive decline, and risk of conversion from MCI to AD^13,14^. Microarray analysis studying 17 brain regions of AD patients in varying disease stages showed gene expression abnormalities of enzymes involved in sphingolipid metabolism, favoring ceramide accumulation and reducing S1P synthesis with disease progression^32^. Analysis of post-mortem AD brains has reported that S1P concentration declined with the progression of AD lesions^30^. This reduction might be caused by the downregulation of sphingosine kinase (SphK) 1 and 2^30,33^, and the upregulation of sphingosine-1-phosphate lyase (S1PL)^32,33^. Decrease in cytoplasmatic SphK2 expression positively correlates with Aβ deposits in AD brains, where SphK2 is preferentially localized in the nucleus, suggesting a disrupted equilibrium between cytosolic and nuclear SphK2^34^. Furthermore, in vitro studies in APP-overexpressed PC12 cells showed that Aβ decreases the expression and activity of SphK1/2 as well as S1P levels^35^. Cerebrospinal fluid (CSF) metabolomic analysis showed elevated S1P levels in MCI subjects and decreased S1P levels in AD patients^36^. However, the role of S1P in AD remains controversial. Inhibition of SphKs activity or overexpression of S1P degrading enzymes showed that S1P stimulates the β-secretase beta-site amyloid precursor protein cleaving enzyme 1 (BACE1) by direct interaction with the enzyme increasing Aβ production in vitro studies^37^. Increased intracellular levels of S1P induced apoptosis in hippocampal and S1PL-deficient neurons^38,39^, and disrupts APP metabolism, which was associated with impaired lysosomal degradation in S1PL-deficient cells^40^.

Recently, in a multi-omics approach^41^, we identified potential targets in the sphingomyelin pathway that suggested modulators of S1P metabolism as possible candidates for AD treatment and tested successfully that treatment with Fingolimod (FTY720), an S1P receptor (S1PR) modulator, was able to recover synaptic plasticity and learning and memory in APP/PS1, an animal model for AD. To further investigate how FTY720 alleviates synaptic plasticity and cognitive impairment, we now explore metabolic abnormalities in APP/PS1 mice, metabolic effects of FTY720 treatment, and their relation to the improvements in synaptic plasticity and memory.

## Results

### APP/PS1 mice exhibit metabolic changes in the brain compared to wild-type mice

Metabolomic data from Biocrates MxP Quant 500 XL assay, together with calculated metabolic indicators, were first analyzed to find differences related to the APP/PS1 genotype. We compared APP/PS1-vehicle and wild-type (WT)-vehicle groups with a series of linear regression models, controlling for sex. We detected several significant (false discovery rate (FDR) ≤ 0.05, or p-value ≤ 0.05 when consistent with another tissue type with FDR ≤ 0.05) changes in the brain tissue, particularly in the parietal, frontal, and temporal regions. Among small molecules, spermidine synthesis (ratio spermidine/putrescine) was downregulated (for illustration: parietal – β (effect size normalized to 1 standard deviation of WT)=-1.78, p=1.2e-5, FDR=3.1e-4; frontal – β=-1.37, p=1.5e-3, FDR=0.034). Other changes were related to lipids: Several lysophospholipid classes were skewed towards lipid species with increased saturated fatty acid (SFA) and decreased unsaturated fatty acid (UFA; consisting of monounsaturated (MUFA) and polyunsaturated (PUFA)) residues, including lysophosphatidylethanolamines (LPEs), lysophosphatidylcholines (LPCs), and LPE plasmalogens. Ratio of phosphatidylinositol (PI) 18:0_20:4 (the most predominant PI) to total PIs was increased. Furthermore, the ratio of odd-chain to even-chain fatty acid sphingomyelins (SMs) was reduced. The tissue-specific effect size and significance are depicted in Figure 1. These metabolic indicators also provided a very high separability between the groups, generally achieving the area under receiver operating characteristic curve (AUC) score over 90% (maximum 96% (95% confidence interval (CI_95_)=90%-100%) for the ratio of MUFA/SFA LPCs in the parietal brain region).

**Fig. 1:**
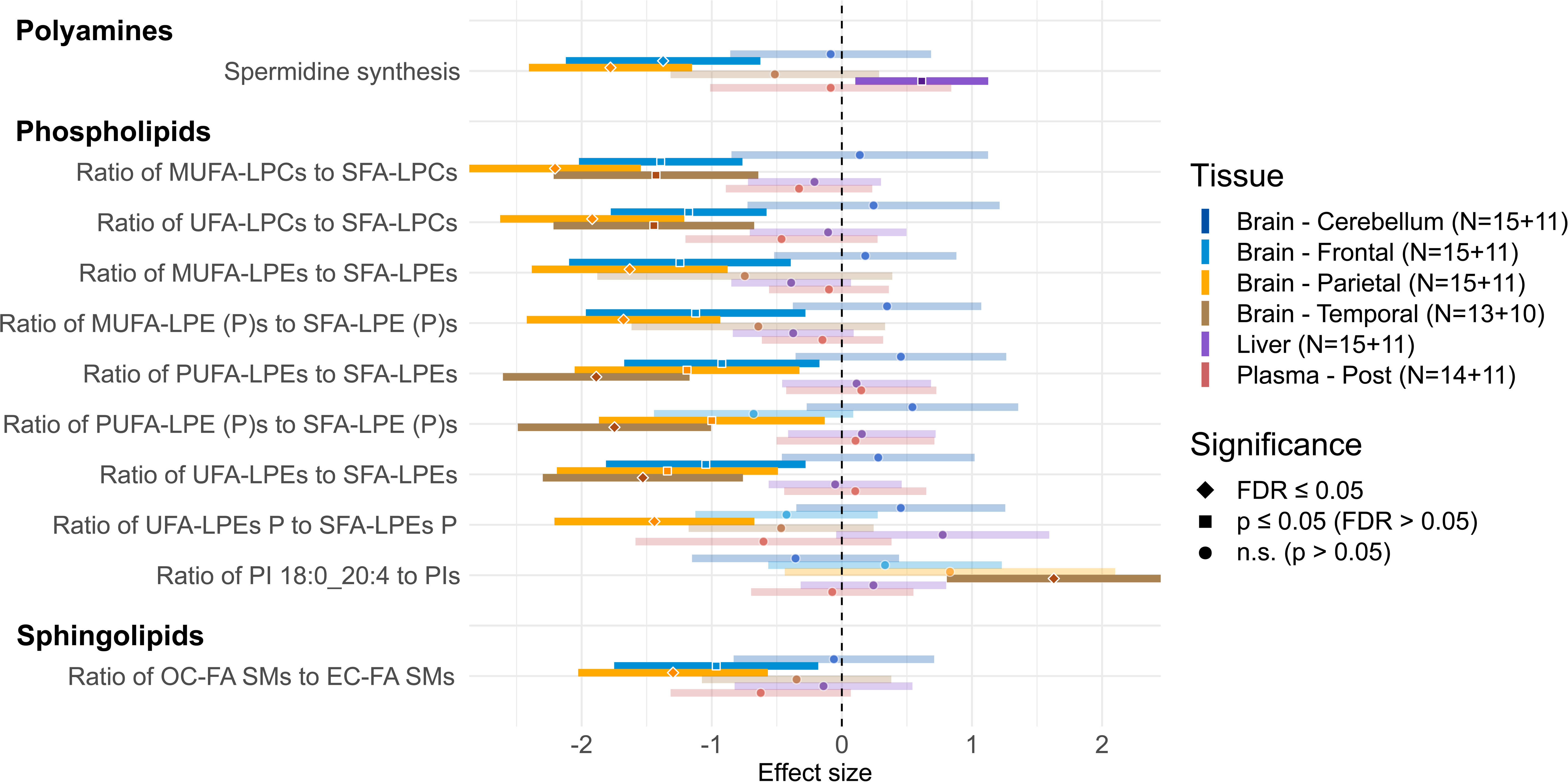
Metabolic alterations in APP/PS1 mice. Forest plot with metabolic differences in APP/PS1-vehicle mice that are statistically significant (FDR ≤ 0.05 for at least 1 tissue) compared to WT-vehicle in a linear regression model (see Methods for details). The normalized regression coefficients are shown on the x axis as points, with the shape denoting the statistical significance (diamond – FDR ≤ 0.05; square – FDR > 0.05 and p ≤ 0.05; circle – p > 0.05), located in the middle of a horizontal line depicting 95% confidence intervals, with lower opacity for insignificant results (p > 0.05). The vertical dashed line across 0 represents no group difference. Positive values mean an increase in APP/PS1-vehicle group. Individual types of tissue are color-coded (dark blue – cerebellum; light blue – frontal cortex; orange – parietal cortex; brown – temporal cortex; purple – liver; red – plasma). Sample counts for each tissue and group are included as “N” with WT group listed first.

### FTY720 affects multiple areas of metabolism in different tissues in APP/PS1 mice

Next, we explored the overall effects of FTY720 on metabolism in APP/PS1 mice, comparing the APP/PS1-FTY720 and APP/PS1-vehicel groups using the same regression approach. All significant findings are presented in Figure 2 (see Supplementary Figure 1 for individual species of complex lipids). Importantly, we observed changes in the sphingolipid metabolism, which is the area directly impacted by the drug mechanism: FTY720 administration led to an increase in S1P in frontal and parietal cortex, and to decrease in sphingomyelinase activity in frontal and temporal cortex and plasma, resulting in lower total plasma ceramides.

**Fig. 2:**
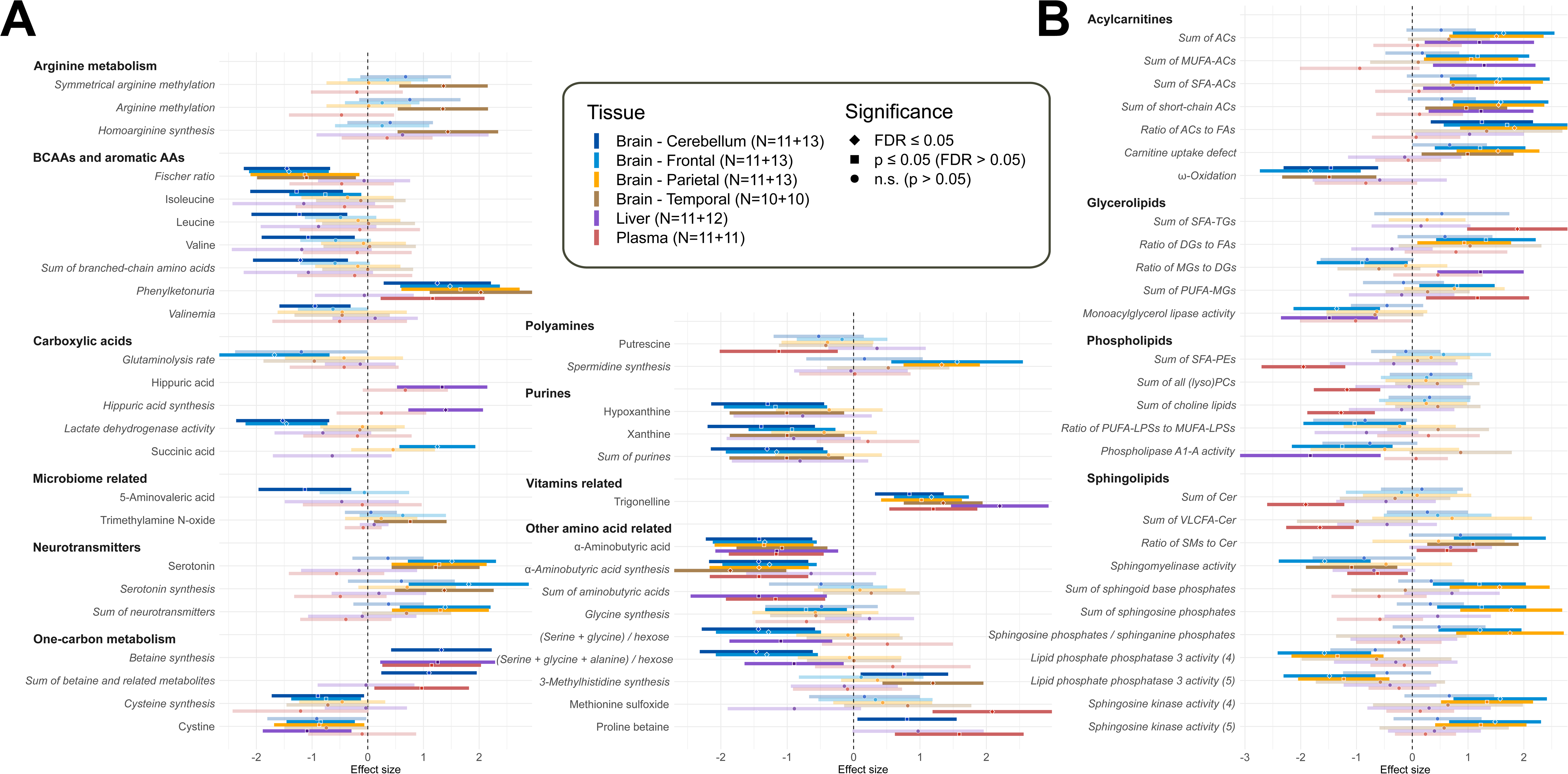
Metabolic effects of FTY720 treatment in APP/PS1 mice. Forest plot with metabolic differences in APP/PS1-FTY720 mice that are statistically significant (FDR ≤ 0.05 for at least 1 tissue among APP/PS1 mice or APP/PS1 and WT combined) compared to APP/PS1-vehicle mice in a linear regression model (see Methods for details). The normalized regression coefficients are shown on the x axis as points, with the shape denoting the statistical significance (diamond – FDR ≤ 0.05; square – FDR > 0.05 and p ≤ 0.05; circle – p > 0.05), located in the middle of a horizontal line depicting 95% confidence intervals, with lower opacity for insignificant results (p > 0.05). The vertical dashed line across 0 represents no group difference. Positive values mean an increase in APP/PS1-FTY720 group. Individual types of tissue are color-coded (dark blue – cerebellum; light blue – frontal cortex; orange – parietal cortex; brown – temporal cortex; purple – liver; red – plasma). Sample counts for each tissue and group are included as “N” with APP/PS1-vehicle group listed first. Calculated metabolic indicators are displayed in italics for easier visual separation from measured metabolites. For complex lipids, only metabolic indicators are shown.

Two FTY720-related changes in small molecules were consistent across all analyzed tissues: increase in trigonelline, which is a methylbetaine form of niacin (vitamin B_3_), and decrease in α-aminobutyric acid (AABA), which is produced from α-ketobutyric acid downstream of homocysteine transsulfuration pathway. Additionally, several amino acid indicators were consistently changed across all analyzed brain regions: phenylketonuria test (ratio tyrosine/phenylalanine) was elevated, and Fischer ratio (ratio branched-chain amino acids (BCAAs) / aromatic amino acids) was lower. BCAAs alone were also decreased in cerebellum.

Multiple brain areas were further altered by FTY720 in the following metabolic pathways: neurotransmitters (increased serotonin), transsulfuration pathway (decreased cysteine synthesis and cystine, higher betaine (trimethylglycine) synthesis), polyamines (increased spermidine synthesis), purines (lower xanthine and hypoxanthine), acylcarnitines (ACs; general increase without a similar increase in fatty acids, and decreased indicator of ω-oxidation), histidine metabolism (higher 3-methylhistidine synthesis), and several indicators with sum of hexose sugars (decreased lactate dehydrogenase activity (lactate/hexose), decreased ratio (serine+glycine)/hexose).

Other FTY720-related changes were typically present in a single type of tissue or single brain area, including increase in indicators with arginine in denominator, increase in carboxylic acids (hippuric acid synthesis, succinic acid), increase in amino acid-related compounds (methionine sulfoxide, proline betaine), and changes in lipids suggestive of decreased monoacylglycerol lipase and phospholipase A1-A activity, along with increased choline lipids and SFA PEs.

### FTY720 improves or normalizes most metabolic changes related to APP/PS1 genotype

In the next step, we searched for metabolic changes in APP/PS1-vehicle mice which were substantially reverted (by at least 50% of the effect) in APP/PS1-FTY720 group (Figure 3A) and changes which showed little or no improvement (less than 50% of the effect; Figure 3B). Note that here we used information from all 4 mouse groups, which led to detection of more genotype-related changes than in the previous 2-group comparison presented in Figure 1 (see Methods for details).

**Fig. 3:**
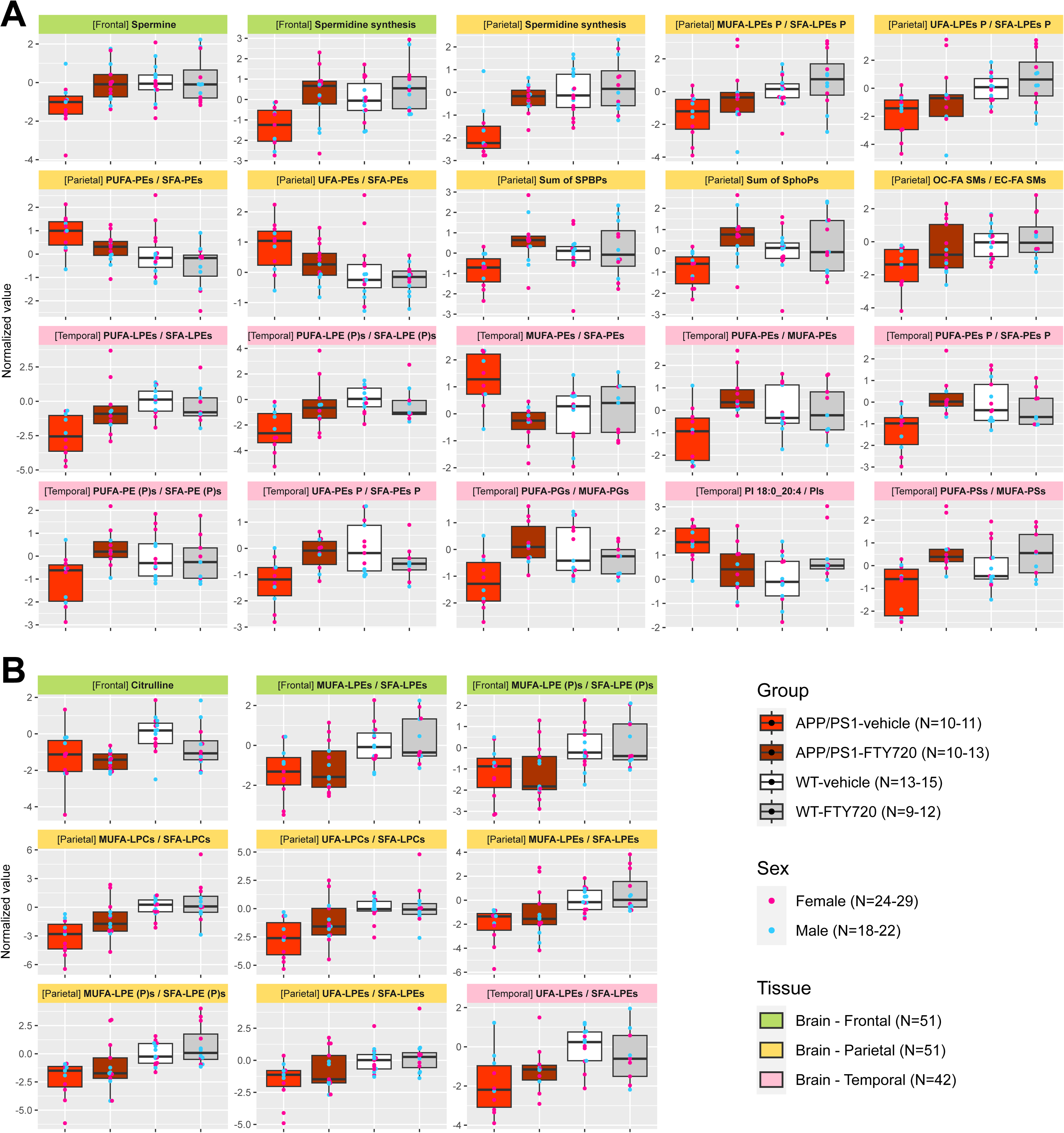
Effect of FTY720 treatment on metabolic alterations in APP/PS1 mice. Overview of analytes differentially present in APP/PS1 mice when considering all groups (see Methods for details) that showed A) strong improvement (≥ 50% of the genotype effect) by FTY720, and B) little to no improvement (< 50% of the genotype effect) by FTY720. The data show normalized values in individual box plots for each of the analytes and the respective tissue (headings color: green – frontal cortex; gold – parietal cortex; pink – temporal cortex). Each box (center line – median; box limits – upper and lower quartiles; whiskers – 1.5× interquartile range) represents one group (bright red – APP/PS1-vehicle; dark red – APP/PS1-FTY720; white – WT-vehicle; gray – WT-FTY720) and is overlayed with points corresponding to individual samples (blue – male; pink – female). Sample counts are included as “N” for each group and each sex with the minimum-maximum range across individual tissues, and for each tissue in total.

Observed changes in spermidine synthesis (frontal and parietal cortex) and spermine (frontal cortex) were completely normalized with FTY720 (Figure 3A). The decreased sum of S1Ps (parietal cortex) was also normalized, trending to even higher concentrations compared to WT. Similarly strong normalization was apparent in ratios with MUFA, PUFA, and SFA phospholipids in temporal cortex (MUFA/SFA and PUFA/MUFA phosphatidylethanolamines (PEs), PUFA/SFA PE plasmalogens, PUFA/MUFA phosphatidylglycerols (PGs), and PUFA/MUFA phosphatidylserines (PSs)). This improvement was only partial in PUFA/SFA PEs in parietal cortex. Several saturation ratios with lysophospholipids were also improved (PUFA/SFA LPEs in temporal cortex, MUFA/SFA LPE plasmalogens in parietal cortex; Figure 3A), while others showed little (MUFA/SFA LPCs in parietal cortex) or no (MUFA/SFA LPEs in frontal and parietal cortex) improvement (Figure 3B). No correction further occurred for decreased citrulline in frontal cortex. On the other hand, the increased ratio PI 18:0_20:4 / total PIs was normalized and the decreased ratio of odd-chain to even-chain fatty acid SMs was partially normalized (Figure 3A).

### FTY720 recovers Barnes memory task and long-term potentiation at the Shaffer collaterals-CA1 (CA3-CA1) synapse

To characterize whether FTY720 treatment can produce recovery in memory, we tested two memory behavioral tasks: the Barnes maze and the novel object recognition (NOR) behavioral tasks. We found that the APP/PS1 group treated with FTY720 was able to recover memory as WT animals in the Barnes maze task (Figure 4: top left). However, when NOR was tested, we found a trend but not significant differences between groups (Figure 4: top center). We then tested whether the treatment could rescue impaired synaptic plasticity in APP/PS1. We measured long-term potentiation (LTP) and post-tetanic potentiation (PTP) in two synaptic inputs: CA3-CA1 and layer II intra-cortical lateral entorhinal cortex (LEC-LEC), located in the hippocampus and in the LEC, respectively. We found a significant recovery of LTP in CA3-CA1 in APP/PS1 animals treated with FTY720 and only a trend in the LEC-LEC LTP (Figure 4: bottom left panels). No differences were found in the PTP measurements (Figure 4: bottom right panels) and no significant differences were found in distance travelled as a measurement of motor skills among groups (Figure 4: top right panel).

**Fig. 4:**
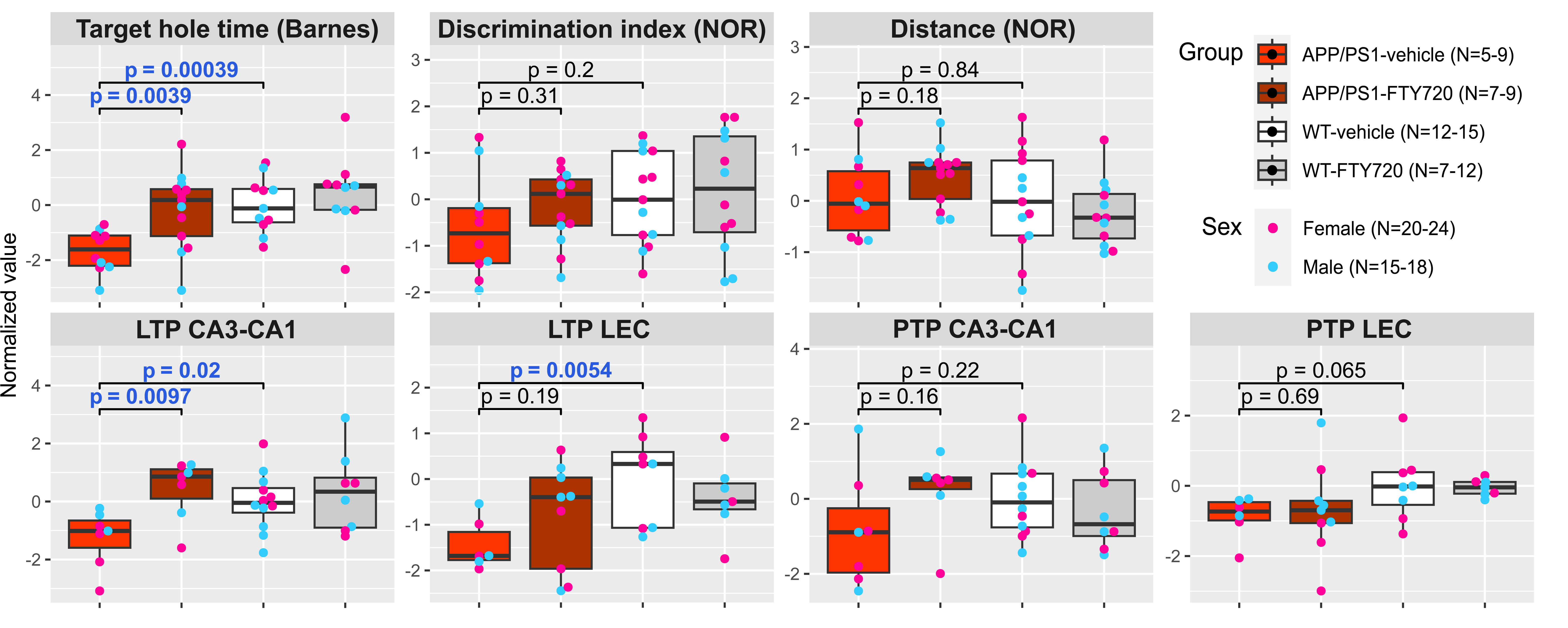
Effect of APP/PS1 genotype and FTY720 treatment on phenotype. Box plots for normalized values of individual behavioral (Target hole time – Barnes maze; Discrimination index and Distance – Novel object recognition test) and electrophysiological (LTP – long-term potentiation; PTP – post-tetanic potentiation; both in either CA3-CA1 hippocampal region or lateral entorhinal cortex (LEC)) measures. Each box (center line – median; box limits – upper and lower quartiles; whiskers – 1.5× interquartile range) represents one group (bright red – APP/PS1-vehicle; dark red – APP/PS1-FTY720; white – WT-vehicle; gray – WT-FTY720) and is overlayed with points corresponding to individual samples (blue – male; pink – female). Welch’s t-test p-values are provided for comparison between WT-vehicle and APP/PS1-vehicle groups, and between APP/PS1-vehicle and APP/PS1-FTY720 groups (p ≤ 0.05 are highlighted with blue font). Sample counts are included as “N” for each group and each sex with the minimum-maximum range across individual measurements.

### Correspondence between metabolic changes and FTY720-related normalization of behavior and neuronal activity in APP/PS1 mice

Finally, we searched for metabolites and metabolic indicators (collectively referred to as analytes) which would best explain the significant improvement and normalization of behavioral and electrophysiological tests in APP/PS1 mice on FTY720, to detect areas of metabolism potentially directly related to the mechanism of the phenotype rescue. The significant normalization was observed for the time to reach the escape hole (target hole time) in Barnes maze and for LTP in CA3-CA1 hippocampal region. Note that these measures are not correlated among APP/PS1 mice (Pearson’s r=-0.09, p=0.77), so both normalizations are likely independent of each other. The 10 best corresponding analytes (see Methods for details on the selection procedure and Supplementary Figure 2 for distribution of selection scores across tissues) for each of the two measures are presented in Figure 5.

**Fig. 5:**
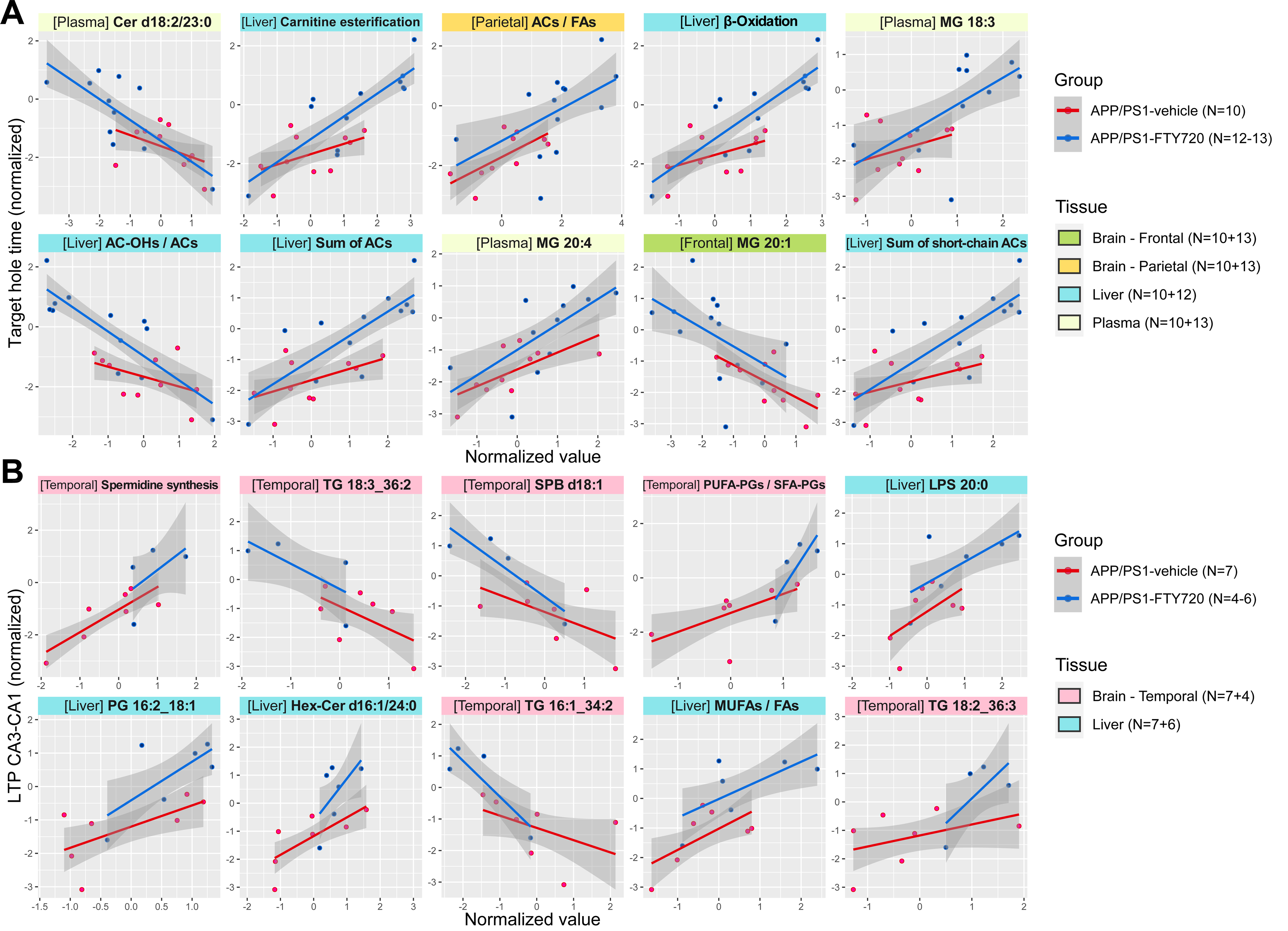
Top metabolic analytes best corresponding to FTY720-related correction of abnormal phenotype in APP/PS1 mice. Scatter plots for top 10 analytes best corresponding (see Methods for details) to A) Target hole time (Barnes maze), and B) LTP in CA3-CA1 hippocampal region, among APP/PS1 mice across tissues. The values of these measurements (y-axis) and the respective top analytes (x-axis) are normalized. The type of tissue is specified and color-coded in the plot headings (green – fontal cortex; gold – parietal cortex; blue – liver; light green – plasma; pink – temporal cortex). Points represent individual samples and lines show the linear trend with 95% confidence intervals as shaded areas, with groups color-coded (red – APP/PS1-vehicle; blue – APP/PS1-FTY720). Sample counts are included as “N” for both groups with the minimum-maximum range across individual tissue types, and for each tissue with the number of samples in both groups where APP/PS1-vehicle is listed first.

The improvement in target hole time (Figure 5A) best corresponds to decrease in ceramide d18:2/23:0 in plasma, to increase in acylcarnitines, β-oxidation, and related indicators in liver and parietal cortex, and to increase in monoacylglycerol (MG) 18:3 and MG 20:4 in plasma but decrease in MG 20:1 in frontal cortex. For LTP in CA3-CA1 region (Figure 5B), the improvement best corresponds to changes in temporal cortex: increase in spermidine synthesis, decrease in triglyceride (TG) 18:3_36:2 and TG 16:1_34:2 but increase in TG 18:2_36:3, decrease in sphingosine d18:1, and increase in ratio PUFA/SFA PGs, and further to changes in liver: increase in lysophosphatidylserine (LPS) 20:0, PG 16:2_18:1, hexosylceramide (Hex-Cer) d16:1/24:0, and ratio of free MUFAs to all free fatty acids.

## Discussion

FTY720 is a medication clinically approved for treatment of multiple sclerosis and is also under investigation for possible use in other neurodegenerative diseases due to its direct modulation of sphingolipid metabolism. In this study, we explored the effect of FTY720 on metabolomic, behavioral, and physiological alterations in APP/PS1 mice as a model of AD. We searched for metabolic areas best corresponding to the observed phenotypic improvements and further characterized the overall effect of FTY720 on metabolic processes.

There has been limited FTY720 research in AD models. Two studies using 5xFAD female mice found that FTY720 treatment halted both spatial memory decline assessed in Morris Water Maze (MWM) and expression of pathological biomarkers^42,43^. Both studies had younger mice subjected to FTY720. Another study found brain gene expression changes in FVB-Tg females after 2 weeks of FTY720 treatment^44^. A reversal treatment study found that 8-week FTY720 treatment recovered deficits in dendritic spines, CA3-CA1 synaptic plasticity, and spatial memory in MWM in 8-month-old APP/PS1 males^45^. Lastly, in our previous study we expanded previous observations on the positive effects of FTY720 treatment in AD models and examined more compromised 9-month-old APP/PS1 mice, and unlike previous studies, administered the drug in water instead of IP injection, performing NOR task, Barnes maze, and synaptic plasticity measurements (CA3-CA1, LECII-LECII)^41^. Here, we replicated the FTY720 effects in Barnes maze and CA3-CA1 LTP but did not find significant differences when performing NOR and LEC-LEC LTP although the trends were apparent. We attribute these results to the smaller number of animals used per group and the robustness of Barnes and conventional CA3-CA1 LTP compared to NOR and LEC-LEC LTP.

The behavioral and physiological improvements by FTY720 were reflected in the metabolomic signature of APP/PS1 mice as highlighted in Figure 6, which integrates pathways with significant drug-related normalization and top correspondence to the functional improvement. The detection of low S1P in parietal cortex (and similar trend in frontal cortex) of APP/PS1 mice is consistent with a report of their decreased neuronal sphingosine kinase concentrations^41,46^. Together with the S1P normalization on FTY720, these results confirm the involvement of the sphingolipid metabolism in the pathology and validity of the initial hypothesis of FTY720 as a suitable modulator that can repair the “sphingolipid rheostat” – recover the balance between pro-apoptotic and pro-survival sphingolipid signals^23^. This is underlined by the decreased indicator of sphingomyelinase activity in frontal and temporal cortex and plasma, resulting in significantly lower plasma ceramides, another anti-inflammatory and pro-survival change^23^. It has been shown that exposure to Aβ peptide 42 (Aβ_42_), which is involved in AD^47^, produces mitochondrial dysregulation leading to a hypometabolic state^48^ affecting neurons^49^, endothelial cells^50^ and glia^51,52^. Aβ_42_ activates sphingomyelinase in murine astrocytes, leading to ceramide-mediated suppression of mitochondrial adenosine triphosphate (ATP) release and consequent fragmentation^53^. This is an important mechanistic link between AD pathology, sphingolipid disruption, and mitochondrial damage, along with the target of FTY720 treatment. Both the increase of sphingosine kinase activity and the decrease of sphingomyelinase activity are well-documented effects of FTY720^44,54^. The presence of ceramide (Cer d18:2/23:0) as well as sphingosine (SPB d18:1), which has been shown to disturb mitochondrial membrane potential in rat hippocampus^55^, among the analytes with top correspondence to FTY720-related behavioral and physiological improvements strongly suggests that reduction of these pro-inflammatory and pro-apoptotic sphingolipids may be the driving factor in the AD phenotype normalization.

**Fig. 6:**
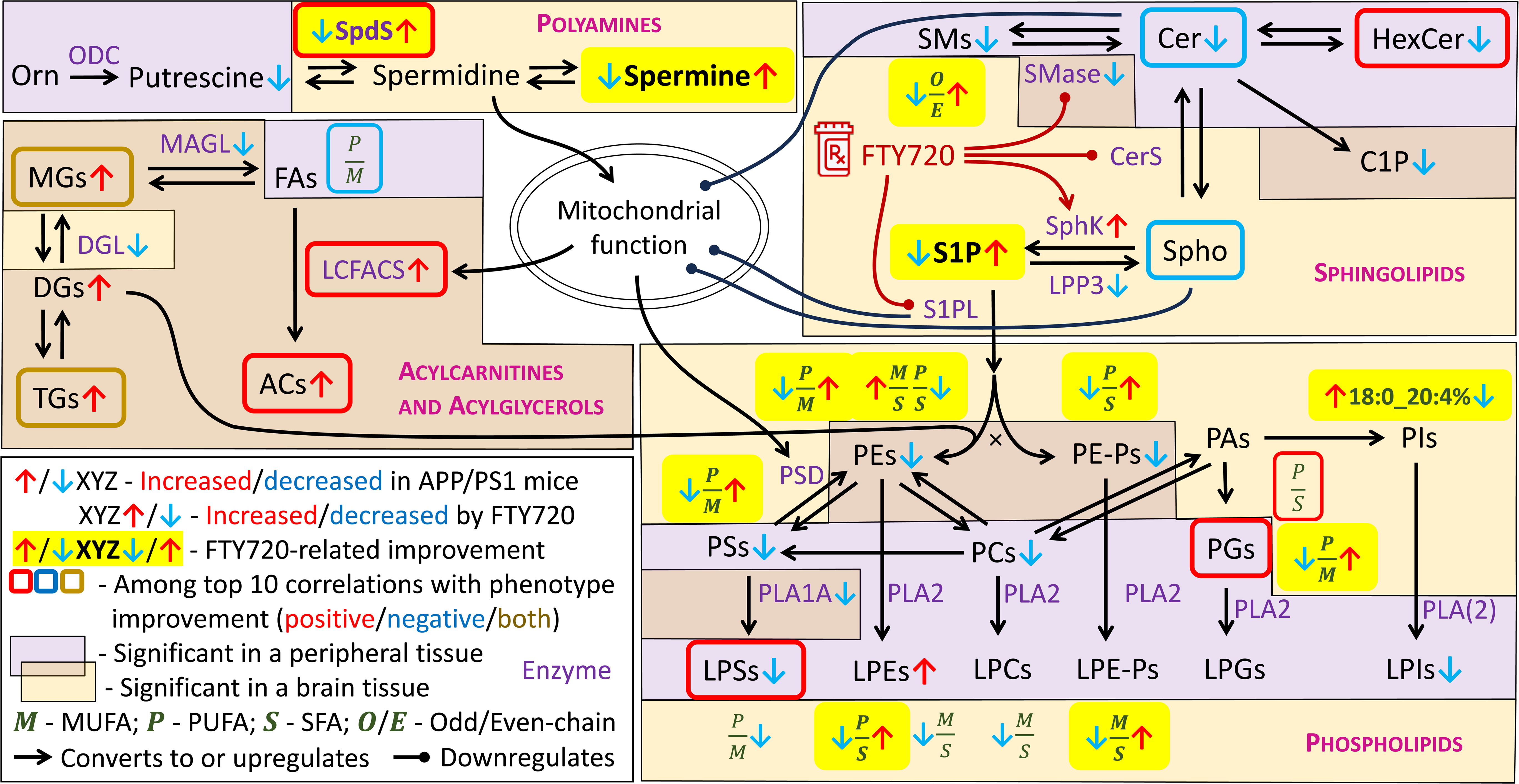
Metabolic areas involved in FTY720-related rescue of APP/PS1 changes in metabolism and phenotype. Pathway map encompassing APP/PS1 metabolic alterations corrected by FTY720 and top 10 metabolic analytes best corresponding to FTY720-related correction of abnormal behavioral and electrophysiological tests in APP/PS1 mice. The known actions of FTY720 and relationships with mitochondrial function are included. The conversion between metabolites (black font) or upregulation are denoted by a regular black arrow, sometimes accompanied by the respective enzyme name (purple font; corresponds to metabolic indicators calculated as ratios from the main measured substrate and product metabolites), whereas downregulation is symbolized by a round-headed curve. For visual clarity, not all existing reactions and connections are shown. Ratios of lipid subgroups (green font in italics) are positioned adjacent to the respective lipid class. Mutually exclusive pathway split is marked with black “×”. Observed changes (red arrow up – increase; blue arrow down – decrease) are distinguished as related to APP/PS1 genotype (arrow before the label) and FTY720 action (arrow after the label), further highlighted where FTY720 correction occurred (yellow background). Top analytes best corresponding to phenotypic improvements are framed (red – positive correlation; blue – negative correlation; dark yellow – group contains metabolites correlated in both directions). The presence of the changes is also divided into central system (brain; light ochre background) and peripheral system (plasma and liver; light purple background), with possible overlap.

Another area of metabolism that we observed altered in brain of APP/PS1 mice are polyamines, specifically lower spermidine synthesis (with spermidine itself also lower based on p-value but with higher FDR) and spermine. These results are inconsistent with an existing report of minimal differences in spermine and spermidine in frontal cortex of APP/PS1 mice of the same age (9 months) and significant elevation at 17 months^56^. Another study found an increase in spermine and spermidine in brain of APP/PS1 mice at 8 months but no difference before or after that up to 18 months although age-related fluctuations were obvious^57^. These findings cannot be directly reconciled and may be affected by the resolution of the brain area location and the exact mouse age. In this study, we observed complete normalization of polyamine values on FTY720 in frontal and parietal cortex of APP/PS1 mice and the indicator of spermidine synthesis in temporal cortex became the top analyte best corresponding to LTP in CA3-CA1 hippocampal region in APP/PS1 mice, suggesting a strong role in the rescue of neuronal function. Declarative memory requires the interaction between hippocampus and surrounded cortical areas including temporal cortex^58^. The hippocampus is instrumental in the initial learning and encoding of experiences, whereas medial temporal lobe cortex is required for retrieval of remote memories^59^. Our observations in the Barnes maze thus indicate that FTY720 affects this memory system through normalization of polyamines. The physiological functions of spermidine and spermine are numerous and diverse, ranging from free radical scavenging and ion channel modulation to autophagy induction and cellular growth regulation^60^. Due to these far-reaching effects, concentrations of spermidine and spermine are tightly regulated and their disruption leads to pathological states^60^. Spermidine also promotes mitochondrial respiration via hypusination of eIF5A (eukaryotic translation initiation factor 5A), directly influencing β-oxidation energy metabolism^61,62^. The lower rate of spermidine synthesis that we detected in brain of APP/PS1 mice thus represents another impact on mitochondrial function and could act synergistically alongside the sphingolipid disbalance. Consistent with the improvement of mitochondrial β-oxidation by FTY720, we observed a drug-related increase across a range of acylcarnitines, which are the β-oxidation intermediates. The sum of acylcarnitines further correlated well with spermidine concentrations across multiple brain regions and plasma (parietal – Pearson’s r=0.52, p=7e-5; frontal – r=0.51, p=1e-4; cerebellum – r=0.44, p=0.001; plasma – r=0.37, p=0.012), supporting the known causal relationship. Moreover, multiple acylcarnitine-related indicators, mainly in liver but also in parietal cortex, ranked among the top analytes best corresponding to the target hole time behavioral score, further supporting the importance of this pathway in the APP-PS1 phenotype improvement with FTY720.

Most of the observed changes in APP/PS1 mice, however, were related to ratios of different fatty acid saturation subgroups in multiple phospholipid classes in brain. While these ratios are rarely investigated, alteration among phospholipids or saturation subgroups of free fatty acids are frequently listed for APP/PS1 mice in a review of lipidomic studies across AD mouse models^63^. In concordance with our results, the most often detected changes include (L)PEs and (L)PCs, which are the most abundant phospholipid classes in neuronal cells^64^, and alterations in other types of phospholipids such as PSs, PIs, and PGs are also reported^63^. The observed direction of lower ratios of UFA to SFA and PUFA to MUFA lipid species in APP/PS1 mice is consistent with known effects of dietary fat on cognition in human where SFAs generally increase the risk of dementia while UFAs, especially ω-3 PUFAs, are protective^65^. The only exception that we detected was for PEs, where the ratio of UFAs to SFAs was increased. A closer look reveals that the decreased ratio of PUFA to MUFA lipids was detected for PEs, PSs, and PGs, which are all mainly localized at mitochondrial membranes and mitochondria-associated membranes of endoplasmic reticulum^66^, suggesting a single mutual point of the disruption. PEs are synthesized in mitochondria from PSs through the action of phosphatidylserine decarboxylase (PSD) although there is another major pathway for PE synthesis from cytidine diphosphate (CDP)-ethanolamine and diglycerides (DGs). CDP-ethanolamine is generated from phosphoethanolamine, one of which sources is S1P through the action of S1PL. Interestingly, the CDP-ethanolamine pathway generates PEs with mainly MUFAs or diunsaturated PUFAs connected to the middle carbon of the glycerol backbone, while the mitochondrial PSD pathway produces higher-order PUFA chains there^67^. Furthermore, both pathways are co-regulated to maintain a stable concentration of PEs^67^. Decreased ratio of PUFA/MUFA PEs in APP/PS1 mice, therefore, implies downregulated mitochondrial PSD pathway, which should be accompanied by increase in CDP-ethanolamine pathway. This is consistent with the observed increase in the ratio of MUFA/SFA PEs and not PUFA/SFA PEs. Note that the presented increase in ratio of PUFA/SFA PEs was detected in a different brain region (parietal cortex) than the other changes in phospholipids discussed so far (temporal cortex), suggesting spatial specificity of the disruptions. The shift in the balance of PE synthesis pathways is then likely further reflected in the observed altered composition of LPEs and (L)PE plasmalogens, as they partake in the PE regulatory mechanism^67,68^. However, it is not clear whether the primary disruption is initiated by reduced flow through the PSD pathway or increased flow through the CDP-ethanolamine pathway. We have already discussed several ways how the observed disturbances in polyamines and sphingolipids could negatively affect mitochondrial function in APP/PS1 mice, which is further supported by a report of reduced mitochondrial mass in this strain^69^. Mitochondrial insufficiency could result in the need for higher CDP-ethanolamine synthesis and potentially even increased S1PL activity, thereby S1P reduction. Alternatively, a priori increased S1PL activity could upregulate the CDP-ethanolamine pathway by making its intermediate phosphoethanolamine more available for conversion into CDP-ethanolamine and the PSD pathway would be downregulated as a secondary event. Indeed, S1P has been shown to interfere with PSD in glioma cells^70^. In either case, increased S1PL activity has been documented in human AD brains and correlated with Aβ deposits^33^, and moreover, S1PL byproduct 2-hexadecenal exhibits pro-apoptotic effects and can lead to decrease in mitochondrial membrane potential^71^. This change is opposed by FTY720, which has been shown to decrease S1PL activity, apparently through competitive inhibition^72^. Importantly, FTY720 normalized most of the phospholipid changes observed in APP/PS1 mice in our experiment, especially in the temporal cortex, suggesting that the balance in pathways for PE synthesis was restored. More research would be needed to confirm whether the normalization was done primarily through decreasing S1PL activity and CDP-ethanolamine pathway, or through improvement in mitochondrial function and restoring PSD activity. None of the PE-related changes ranked among the top analytes best corresponding to the behavioral and electrophysiological improvement, although for other phospholipids, the ratio of PUFA/SFA PGs in temporal cortex showed a high positive correlation with LTP CA3-CA1. This might reflect a variation in mitochondrial structural properties as PGs are precursors for the synthesis of cardiolipins, important constituents of mitochondrial membranes^66^.

Besides the correction of APP/PS1-related metabolic changes, FTY720 affected multiple other parts of metabolism. To help explain these observations, we integrated current knowledge about signaling and pathways known to be altered by FTY720 (Figure 7). Unlike the enzymes involved in sphingolipid metabolism, many of the effects on signaling are shared by FTY720 and S1P due to their structural similarity, action on S1PRs, or elevation of S1P by FTY720. The most explored pathways affected by FTY720 and S1P are cyclic adenosine monophosphate (cAMP) / cAMP response element-binding protein (CREB)^73–75^, and brain-derived neurotrophic factor (BDNF) / tropomyosin receptor kinase B axis^74–76^, amplified by histone deacetylases^77–79^, and with a downstream effect on phosphatidylinositol 3-kinases (PI3K) / protein kinase B (Akt) signaling^80^, and Janus kinase (JAK) / signal transducer and activator of transcription (STAT) signaling^81^, both also activating the pro-survival protein myeloid cell leukemia-1 (Mcl-1)^82,83^. BDNF is a critical factor in synaptic plasticity including LTP in hippocampus^84^, consistent with the observed CA3-CA1 LTP normalization by FTY720. BDNF also interacts with the serotonin system, promoting survival and differentiation of serotonergic neurons^85^, which can explain the observed serotonin increase by FTY720. PI3K/Akt and JAK/STAT pathways convey a neuroprotective effect by FTY720 or S1P, including improvement in mitochondrial function and modulation of microglia to pro-survival M2 polarization^86–89^. Mcl-1 has a crucial impact on mitochondrial respiration and energy production, maintaining neuronal survival^90,91^. The cAMP/CREB signaling via S1PRs also activates the master regulator of mitochondrial biogenesis peroxisome proliferator-activated receptor γ (PPARγ) coactivator 1α (PGC-1α)^73^, and additionally, S1P directly binds to and stimulates activation of PPARγ^92^, which promotes the anti-inflammatory M2 microglial polarization^93^. The overall improvement of mitochondrial function and survival could be reflected in the observed increase in acylcarnitines as mediators of mitochondrial fatty acid β-oxidation, and possibly also decreased BCAAs and AABA, since BCAAs are metabolized in mitochondria by branched-chain α-ketoacid dehydrogenase complex (BCKDH) and the same applies for α-ketobutyric acid with BCKDH as an alternative pathway to its metabolism to AABA. Furthermore, both PI3K/Akt and JAK/STAT pathways can activate xanthine oxidoreductase^94^, which might explain the observed decrease in xanthine and hypoxanthine.

**Fig. 7:**
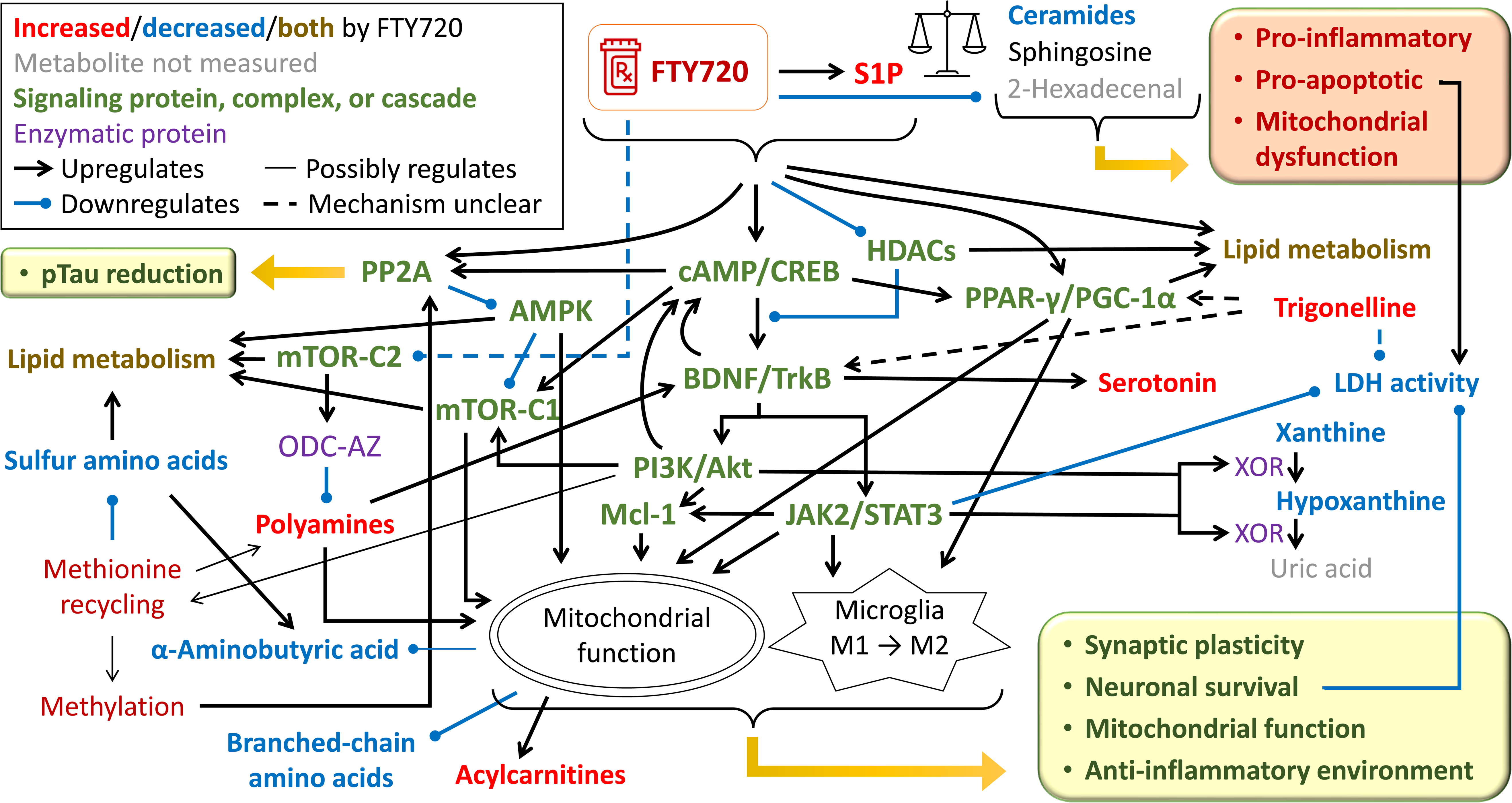
Assumed FTY720 mechanism of action and its effect on metabolism in APP/PS1 mice. Pathway map integrating current knowledge of FTY720 mechanism of action and how its downstream activation of several signaling pathways explains most of the observed metabolic changes related to FTY720 in APP/PS1 mice. The conversion between metabolites, signaling, or upregulation are denoted by a regular black arrow, sometimes accompanied by the respective enzyme name (purple font), whereas downregulation is symbolized by a round-headed blue curve. Relationships with mitochondrial function and switch in microglia polarization are included. The connections are dashed when the mechanism of action is unclear. Thin-line connections represent uncertainty in the strength and quantitative importance of the connection effect. For visual clarity, not all existing reactions and connections are shown. Connections with signaling proteins, protein complexes, and cascades (green font) are based on available literature. Metabolites and metabolic areas are color-coded with respect to the direction of observed change by FTY720 and significance (red – increased by FTY720; blue – decreased by FTY720; dark yellow – members of the class both increased and decreased by FTY720; black – no significant difference (FDR > 0.05); gray – metabolite not measured). For methionine recycling and methylation (dark red), indirect evidence suggests upregulation. The balance icon symbolizes rebalancing the “sphingolipid rheostat”.

The levels of cAMP together with its downstream PI3K/Akt signaling further contribute to activation of mammalian target of rapamycin (mTOR) complex 1 (mTOR-C1)^95,96^, which also stimulates mitochondrial function^97^. Accordingly, increased mTOR activation (or correction of its low activation) has been observed for both S1P and FTY720 although other pathways may have been involved^98,99^. However, mTOR complex 2 (mTOR-C2) is inhibited by FTY720 due to downregulation of its coactivator rapamycin-insensitive companion of mammalian target of rapamycin (RICTOR)^100^, which partakes in mTOR-C2 but not mTOR-C1, although the mechanism behind RICTOR downregulation by FTY720 remains unclear. Interestingly, mTOR-C2 is necessary for synthesis of ornithine decarboxylase (ODC) antizyme (ODC-AZ)^101^, which negatively regulates ODC, one of two rate-limiting enzymes in polyamines synthesis. Downregulation of mTOR-C2 by FTY720 could thus lead to decreased ODC-AZ synthesis and increased ODC activity, generating more polyamines as observed. Besides improving mitochondrial function, spermidine and spermine also increase levels of neurotrophic factors, including BDNF^102^. The other key enzyme for polyamine synthesis is S-adenosylmethionine decarboxylase, which converts S-adenosylmethionine (SAM) to S-adenosylmethioninamine, the required substrate for synthesis of spermidine and spermine. SAM is a universal methyl donor and a part of the methionine cycle (methionine – SAM – S-adenosylhomocysteine – homocysteine (Hcy) – methionine), which can be metabolically exited by SAM decarboxylation, or predominantly, by Hcy transsulfuration. We observed that FTY720 treatment lead to decrease in several metabolites and indicators in the transsulfuration pathway (related to cysteine, cystine, and AABA, while Hcy was not well detected in brain tissue). On the other hand, we saw an increase in many methylated compounds or related indicators in some tissues or plasma (glycine betaine (trimethylglycine), proline betaine (dimethylproline), 3-methylhistidine, trimethylamine N-oxide, trigonelline (1-methylnicotinate), arginine methylation, methionine sulfoxide). Based on these results, it is conceivable that FTY720 favors formation of methionine cycle by Hcy re-methylation rather than metabolism of Hcy through the transsulfuration pathway. This would lead to increased SAM and its decarboxylation to stimulate synthesis of spermidine and spermine. This upregulation of methionine cycle might result from increased PI3K, as it can mediate activation of methionine synthase (MTR)^103^, which re-methylates Hcy to methionine, and methionine adenosyltransferase^104^, which converts methionine to SAM. Previously, we have shown that impaired methionine cycle at methylenetetrahydrofolate reductase (MTHFR), the rate-limiting enzyme producing 5-methyltetrahydrofolate, which is the methyl donor for MTR, leads to downregulation of protein phosphatase A2 (PP2A) and PP2A-mediated tau protein dephosphorylation, effectively resulting in tau hyperphosphorylation^105^, the histopathological hallmark of AD that is thought to follow Aβ accumulation and also occurs in APP/PS1 mice^106^. The involvement of PP2A in AD is underlined by its interaction with apolipoprotein E (ApoE), the most common AD genetic risk factor, where the ApoE risk allele ε4 is associated with decreased PP2A activity^107^ and protective allele ε2 is associated with increased PP2A activity in correlation with tau^108^. Correspondingly, FTY720 and S1P increase PP2A activation although the main mechanism may involve direct binding to PP2A^109^, and additionally, downstream regulation of cAMP signaling^110^. PP2A also further stimulates mTOR-C1 by downregulating its inhibitor adenosine monophosphate-activated protein kinase (AMPK)^111^.

Furthermore, trigonelline, one of the elevated methylated compounds, has been reported to exhibit neuroprotective and anti-diabetic effects via BDNF elevation^112^ and PI3K/Akt signaling^113^, and PPARγ^114^, i.e. similarly to S1P and FTY720, although the exact mechanism has not been elucidated. Trigonelline also reduces lactate dehydrogenase activity (LDH)^115^, again in concordance with effects of FTY720^116^ and S1P through JAK/STAT signaling^117^. Even though these sources measure LDH activity in extracellular fluid or culture media to quantify cellular death, as LDH is released from damaged cells, these relationships are consistent with the observed reduction in the indicator of LDH activity (combining both extracellular and intracellular sources) by FTY720 in several brain regions, along with changes in other indicators related to glucose and lactate.

Overall, the observed metabolic effects of FTY720 are consistent with its known action on several signaling pathways that converge at improving mitochondrial function and increasing pro-survival and anti-inflammatory environment, while FTY720 metabolically inhibits the opposite signals through sphingolipid regulation (Figure 7). Multiple of the involved signaling pathways also affect lipid metabolism, regulating lipogenesis, lipolysis and β-oxidation, and MUFA to SFA conversion^118–120^.

Lastly, we observed consistent FTY720-related elevation of the tyrosine/phenylalanine ratio, which semantically traces phenylalanine hydroxylase (PAH) activity. PAH is a liver enzyme, and we observed the indicator increased in all brain tissues and plasma but not liver, so this disturbance is not related to PAH. Indeed, the change in the ratio is driven exclusively by tyrosine and not phenylalanine. Measured catecholamines were not significantly impacted, so remaining possible explanations include downregulation of tyrosine aminotrasferase, without a clear FTY720-related mechanism, or downregulation of thyroglobulin synthesis in thyroid, which could be contributed by FTY720 anti-inflammatory properties.

In conclusion, we have shown that recovery of hippocampal LTP and Barnes maze performance by FTY720 is accompanied by correction of brain metabolism in polyamines, sphingolipids, and saturation ratios of phospholipids. These changes indicate rebalancing the “sphingolipid rheostat”, reactivating PE synthesis via mitochondrial PSD pathway, and recovering normal polyamine levels supporting mitochondrial activity. The results are consistent with FTY720 promoting mitochondrial function, synaptic plasticity, and anti-inflammatory environment, while reducing pro-apoptotic and pro-inflammatory signals. This study also has several limitations: Not all the analyzed mice underwent the behavioral and physiological testing, which results in higher uncertainty in the correlation estimates. Second, the flow-injection analysis of complex lipids does not always allow to reliably identify the exact forms (e.g. two of the three TG chains are aggregated), and furthermore, the lipid identification specified by the kit manufacturer represents the most likely form among possible isobaric and isomeric species.

## Methods

### Mouse model

All animal procedures and experiments were performed in accordance with the National guidelines (National Institute of Health) and approved by the Institutional Animal Care and Use Committee of the State University of New York, Downstate Health Sciences University. B6.Cg-Tg(APPswe,PSEN1dE9)85Dbo/Mmjax (APP/PS1) mice and C57Bl/6J (WT) mice were purchased from The Jackson Laboratory. The APP/PS1 is a double transgenic mouse expressing a chimeric mouse/human amyloid precursor protein (Mo/HuAPP695swe) and a mutant human presenilin 1 (PS1-dE9) both directed to central nervous system neurons^121^.

### Drug treatment and experimental design

APP/PS1, and their wild-type littermates were used to examine whether FTY720 could ameliorate AD pathology and cognitive deficits in vivo. FTY720 was dissolved in water at a concentration of 10 mg/mL and was given to 7 month old mice through the drinking water at 1 mg/kg/day for four weeks as previously described^41,43^. Drinking water alone was given to mice serving as a vehicle-treated control group. Dark water bottles were used in all groups since the FTY720 is light-sensitive. After FTY720 treatment, mice were subjected to a behavioral battery (Novel Object Recognition Task and Barnes Maze Test). Plasma collection was performed before euthanasia obtained from blood collected from the submandibular vein, then the animals were sacrificed to perform electrophysiological recordings and tissue sample collection of liver, frontal, parietal, and temporal brain cortex, and cerebellum, which were frozen in liquid nitrogen, and stored at –80°C. The complete experimental timeline is depicted in Supplementary Figure 3.

### In vitro electrophysiological recordings

Mice were anesthetized with Ketamine/Xylazine (100/10 mg/kg) and decapitated with an animal guillotine. Horizontal hippocampal slices (400 μm) were prepared using a Vibrotome slicer (VT 1000S; Leica) in ice-cold cutting solution containing the following in mM: 130 potassium gluconate, 5 KCl, 20 HEPES acid, 25 glucose, 0.05 kynurenic acid, 0.05 EGTA-K, and pH equilibrated at 7.4 with KOH. After slicing, the tissue was allowed to recover for an hour before the beginning of experiments in artificial CSF (aCSF) that contained the following in mM: 157 Na+, 136 Cl−, 2.5 K+, 1.6Mg2+, 2 Ca2+, 26 HCO3−, and 11 D-glucose. LTP recordings were performed in an interface chamber (Fine Scientific Tools, Vancouver, Canada) and slices were perfused with aCSF continuously bubbled with 95% O2/5% CO2, to maintain pH near 7.4 and the temperature was set at 34 °C.

Field excitatory post-synaptic potentials (fEPSPs) were recorded in the CA1 stratum radiatum and lateral entorhinal cortex superficial layer II (LECII) with a glass electrode filled with aCSF (2–3MΩ resistance), and the fEPSPs were elicited by stimulating the Schaffer collateral fibers and LECII with a bipolar electrode. Input–output curves were obtained, and a stimulus that evoked ∼40% of the maximum fEPSP was selected to record the baseline. Baseline responses were obtained (15 min with an inter-stimulus interval of 20 s) before high-frequency stimulation (HFS) one train of 100 stimuli at 100 Hz and three trains of 100 stimuli at 100 Hz, with 10 s intervals were used to induce synaptic LTP at the CA3–CA1 and LECII–LECII synapses, respectively. Responses were recorded for 60 min after HFS. The tungsten stimulating electrodes were connected to a stimulus isolation unit (Grass S88), and the recordings were made using an Axoclamp 2B amplifier (Molecular Devices) and then filtered (0.1 Hz–10 kHz using −6 dB/octave). The voltage signals were digitized and stored on a PC using a DigiData 1200A (Molecular Devices) for offline analysis. The fEPSP slope was measured and expressed as a percentage of the baseline. The data was analyzed using Axon pCLAMP software, and the results are expressed as the mean ± standard error of the mean (SEM). Post-tetanic potentiation (PTP) is defined as the average slope of the EPSP during the first minute after high frequency stimulation (HFS). LTP CA3-CA1 or LTP of lateral entorhinal intracortical synapses (LEC-LEC) are defined as the average slope last in the last 5 min (70-75 min) after HFS^41^.

### Novel object recognition (NOR)

Mice were habituated to experimental apparatus consisting of a gray rectangular open field (60 cm × 50 cm × 26 cm) for 5 min in the absence of any objects for 3 days. On the third day, after the habituation trial, mice were placed in the experimental apparatus in the presence of two identical objects and allowed to explore them for 5 min. After a retention interval of 24 h, mice were placed again in the apparatus, where one of the objects was replaced by a novel object. All sessions were recorded using Noldus Media Recorder software. Exploration of the objects was defined as the mice facing and sniffing the objects within 2 cm distance and/or touching them, assessed with ANY-maze software. The ability of the mouse to recognize the novel object (discrimination index) was determined by dividing the difference between exploration time devoted to the novel object and the time devoted to the familiar object by the total time exploring the novel and familiar objects during the test session.

### Barnes maze

The behavioral apparatus consisted of a white flat, circular disk with 20 holes around its perimeter. One hole held the entrance to a darkened escape box not visible from the surface of the board, allowing the subject to exit the maze. The escape chamber position remained fixed during all trials. Mice learn the location of the escape hole using spatial reference points that were fixed in relation to the maze (extra-maze cues). The task consisted of one habituation trial on day 1 where the escape hole was presented to the animal, the animal remained in the escape box for 2 min. After the habituation trial, the training phase consisted of four 3-min trials of spatial acquisition for 4 consecutive days with a 15 min inter-trial interval. On the fifth day (probe trial) the escape box was removed, and the animals were allowed to explore the maze for 90 s. All sessions were recorded using Debut video software and assessed through ANY-maze software. For each trial, several parameters were recorded to assess performance. These include the latency to locate the escape box, the number of incorrect holes checked prior to entering the escape box, as well as the distance traveled prior to locating the escape box. For the probe trial, the time spent on the target hole was analyzed.

### Chromatography and mass spectrometry

Targeted metabolomic analysis was based on triple quadrupole ultra-high-performance liquid chromatography tandem mass spectrometry (UHPLC-MS/MS) using Shimadzu Nexera chromatography platform (Shimadzu Corporation, Kyoto, Japan) coupled to Sciex QTrap 5500 mass spectrometer (AB Sciex LLC, Framingham, Massachusetts, USA). We applied the Biocrates MxP Quant 500 XL targeted metabolomic kit (Biocrates Life Sciences AG, Innsbruck, Austria), potentially quantitating 106 small molecules and free fatty acids in chromatography mode and 913 complex lipids in flow-injection mode (FIA-MS/MS), exploring a broad range of metabolic pathways. Annotations for the individual metabolites with identifiers to external databases are provided in Supplementary Data File 1 (the exact Q1/Q3 mass transitions are not included, as Biocrates prefers to keep this information undisclosed as proprietary knowledge).

Additionally, 474 metabolic indicators were calculated from sums or ratios of relevant metabolites according to Biocrates MetaboINDICATOR formulas^122^ (see Supplementary Data File 2). We refer to the whole set of metabolites and metabolic indicators as “analytes”. The indicators can be regarded as physiologically relevant measures and are statistically analyzed separately from metabolites. The indicators denoted as “X synthesis” are computed as a ratio of metabolite X and its main precursors in an attempt to reflect the conversion ratio. Since there are multiple explanations why such an indicator could be altered, the interpretation needs to be done cautiously in context of the individual metabolites.

Brain and liver samples were extracted in plastic vials with isopropanol at volume 3 µl/mg, homogenized with sonicator, and centrifuged for 20 minutes. 10 µl of the clear tissue extracts, or 10 µl of plasma samples directly, were transferred onto a kit plate with pre-injected internal standards and dried down. In brief, the rest of the assay includes derivatization with 5% phenylisothiocyanate in pyridine, ethanol and water (1:1:1), and subsequent extraction with 5 mM ammonium acetate in methanol. Chromatography was done with 0.2% formic acid in acetonitrile (organic mobile phase) and 0.2% formic acid in water (inorganic mobile phase). Flow-injection analysis was performed with methanol and Biocrates MxP Quant 500 additive of undisclosed composition. All solvents used were of LC/UHPLC-MS grade, except for ethanol with the American Chemical Society and United States Pharmacopeia grade.

Sample handling was done on dry ice to avoid multiple freeze-thaw cycles. We randomized the samples across plates, with stratification, already prior to their processing to avoid any accidental bias towards one of the groups. Plates included blanks (phosphate buffered saline for plasma, isopropanol for tissue extracts) to calculate limits of detection, repeats of a kit quality control sample to calculate concentrations and monitor the coefficient of variation (median for analyzed compounds: <10% for UHPLC, <15% for FIA), and kit calibrators for seven-point calibrations of certain compounds.

### Data preprocessing

Peak areas and concentrations: Mass spectrometry signal was acquired in Sciex Analyst v1.6.24 (AB Sciex LLC, Framingham, Massachusetts, USA) and chromatographic peaks were identified, reviewed, and integrated in the in-house software Integrator (patent pending – see Disclosures for details). Areas of metabolite peaks were divided by areas of their respective internal standards. Further processing was done in R v4.3.2^123^ with RStudio v2023.12.0^124^. For most compounds, concentrations were estimated linearly from expected concentrations in the kit quality control samples using their median, which also serves as a plate normalization. Seven-point linear or quadratic calibration was applied where possible.

Limits of detection (LODs): LODs were calculated as 3× median signals in blanks. Metabolites with more than 50% values below LOD in all subgroups were filtered out. Values below LOD in remaining metabolites were not adjusted, since they represent the best relative estimate of the true values.

Calculated analytes: Metabolic indicators were calculated according to Biocrates MetaboINDICATOR formulas^122^. Ratios with zeros were treated as missing values and not included in the analysis. The metabolic indicators were also plate-normalized using quality control samples.

Data transformation: Additional data transformations were applied for metabolomic, behavioral, and electrophysiological data. To better approximate Gaussian distributions, we applied Box-Cox transformation with R package car^125^. Tukey’s fencing^126^ was used to adjust remote outliers (k=3) to protect against skewing the means by extreme values while not reducing the variance greatly compared to outlier removal. Furthermore, the values were standardized with respect to the reference group (WT-vehicle for most comparisons) to facilitate comparison of regression coefficients in the statistical analysis.

Missing data: Not all mouse subjects underwent the behavioral evaluation and electrophysiological tests and not all tissue samples were harvested successfully. These missing cases are not considered in the analysis, so the numbers of mice in each group slightly differs between the tissue types and between different behavioral and electrophysiological tests.

### Statistical analysis

All statistical analyses were performed in R v4.3.2^123^ with RStudio v2023.12.0^124^, using R package ggplot2^127^ for graphical output and InkScape v1.3 for image layout editing for improved readability. Differential analysis: Behavioral and electrophysiological data were compared directly with Welch’s t-test between the WT-vehicle and APP/PS1-vehicle groups, and between APP/PS1-vehicle and APP/PS1-FTY720 groups. To analyze differences in metabolomic data, a series of multivariable linear regression models were employed, each corresponding to a specific analyte. In these models, the values of the analytes served as the dependent variables, while genotype and/or drug administration status were included as key independent variables. Sex was incorporated as an additional covariate to adjust for potential sex-related differences. The direction and magnitude of the differences were assessed through the estimated regression coefficients for genotype and drug administration status, which quantify the relative change in analyte levels, and p-values were used to evaluate the statistical significance of these estimates, providing a robust framework for interpreting the observed differences. Due to standardization, the regression coefficients have a unit of 1 standard deviation of the distribution of the reference group.

FDR control: For each regression coefficient of interest in the metabolomic data, two-tailed p-values across the models were controlled for FDR using the q-value procedure with R package qvalue^128^. The control was done separately for each combination of plate analytical method (UHPLC, FIA) and analyte type (measured metabolites, calculated indicators). Effects with FDR ≤ 0.05 and effects with p-value ≤ 0.05 only when consistent with another tissue type with FDR ≤ 0.05 were considered statistically significant.

Analysis of FTY720-related metabolic improvement and non-improvement: To detect which analytes were altered due to the APP/PS1 genotype and improved by FT720, we applied the following criteria: 1) Either APP/PS1-vehicle group was significantly different from WT-vehicle group using FDR (direct significance), or APP/PS1-vehicle group was significantly different from all other subjects using FDR (to achieve higher statistical power by including all samples) and significantly different from WT-vehicle group using nominal p-value (to guarantee that the difference is not solely due to the other groups), and 2) the size of the drug correction effect (difference between APP/PS1-FTY720 group and APP/PS1-vehicle group) was at least 50% of the genotype effect (difference between WT-vehicle group and APP/PS1-vehicle group), posing a threshold on practical significance. Analogously, the APP/PS1 genotype was not improved by FTY720 when: 1) Either APP/PS1-vehicle group was significantly different from WT-vehicle group using FDR, or all APP/PS1 subjects were significantly different from all WT subjects using FDR and APP/PS1-vehicle group was significantly different from WT-vehicle group using nominal p-value, and 2) the size of the drug correction effect (difference between APP/PS1-FTY720 group and APP/PS1-vehicle group) was less than 50% of the genotype effect (difference between WT-vehicle group and APP/PS1-vehicle group). Note that this approach, which considers all 4 mouse groups, detects more analytes than the initial differential analysis searching for genotype-related changes, which considers only 2 groups.

Integrative analysis: Behavioral and physiological measures that were significantly altered in APP/PS1-vehicle mice and recovered with FTY720 were analyzed together with metabolomic data to reveal analytes that best correspond to the recovery trajectory. The selection was based on Pearson’s correlation coefficient between the respective behavioral or physiological measure and metabolomic data among the APP/PS1 mice. The magnitude of correlation coefficients in the FTY720 and vehicle subgroups were required to be no less than 50% of the magnitude of correlation coefficient among all APP/PS1 mice to assert certain consistency in the relationship in both subgroups as would be expected for a causal relationship. We selected the top corresponding analytes using the final score of the analyte correlation coefficient minus the group effect size obtained from an additional linear regression model with sex as a covariate (ideally, we would like to see zero additional group effect not explained by the association with the behavioral or physiological measure), both normalized to their maximum-minimum range across all analytes in absolute values.

AUC analysis: We performed a simple univariate AUC analysis using R package pROC^129^ to evaluate separation between selected groups. Confidence intervals are computed using the default DeLong method.

## Supporting information

Suuplementary Figure 1

## Acknowledgements

Dedicated to the memory of Dr. Herman Moreno, who was part of the Alzheimer’s Disease Metabolomics Consortium and left the foundations for this manuscript to be developed.

This project was enabled in part by the Alzheimer’s Disease Metabolomics Consortium (ADMC) U01AG061359 and the Alzheimer’s Gut Microbiome Project (AGMP) U19AG063744 and 3U19AG063744-04S1, with additional funding through R01AG046171, RF1AG059093, RF1AG058942, and RF1AG051550 supported by the National Institute on Aging grants awarded to Dr. Kaddurah-Daouk at Duke University in partnership with multiple academic institutions. As such, the investigators within the ADMC and AGMP not listed in this publication’s authors’ list, provided analysis-ready data, but did not participate in designing the study, conducting the analyses or writing of this manuscript. A complete listing of the AD Metabolomics Consortium (ADMC) investigators can be found at: https://sites.duke.edu/adnimetab/team/. A listing of AGMP Investigators can be found at: https://alzheimergut.org/meet-the-team/. Metabolomics data used in preparation of this article were generated by the Alzheimer’s Disease Metabolomics Consortium (ADMC).

## Data availability statement

Metabolomics data will be available via the AD Knowledge Portal hosted by Sage Bionetworks.

---

**Supplementary Fig. 1: Metabolic effects of FTY720 treatment in APP/PS1 mice on individual lipid species**

Forest plot with metabolic differences in APP/PS1-FTY720 mice that are statistically significant (FDR ≤ 0.05 for at least 1 tissue among APP/PS1 mice or APP/PS1 and WT combined) compared to APP/PS1-vehicle mice in a linear regression model (see Methods for details). This supplementary figure shows results for individual lipid species. The normalized regression coefficients are shown on the x axis as points, with the shape denoting the statistical significance (diamond – FDR ≤ 0.05; square – FDR > 0.05 and p ≤ 0.05; circle – p > 0.05), located in the middle of a horizontal line depicting 95% confidence intervals, with lower opacity for insignificant results (p > 0.05). The vertical dashed line across 0 represents no group difference. Positive values mean an increase in APP/PS1-FTY720 group. Individual types of tissue are color-coded (dark blue – cerebellum; light blue – frontal cortex; orange – parietal cortex; brown – temporal cortex; purple – liver; red – plasma). Sample counts for each tissue and group are included as “N” with APP/PS1-vehicle group listed first.

**Supplementary Fig. 2: Selection of top metabolic analytes best corresponding to FTY720-related correction of abnormal behavioral and electrophysiological tests in APP/PS1 mice**

Scatter plots with distribution of components of the selection score (x-axis – magnitude of Pearson’s correlation coefficient; y-axis – magnitude of normalized group effect from linear regression model; see Methods for details) for A) Target hole time (Barnes maze), and B) LTP in CA3-CA1 hippocampal region, among APP/PS1 mice across tissues. Each point represents one analyte, with the top 5 labeled by text, top 20 showed as full points, and others depicted as circles. Individual types of tissue are color-coded (red – cerebellum; dark yellow – frontal cortex; green – liver; blue– parietal cortex; purple – plasma; pink – temporal cortex).

**Supplementary Fig. 3: Experimental Timeline**

7-month-old animals (APP/PS1 or wild-type littermates) were treated with/without FTY720 for two months. At 9 months they were trained on the NOR (four days) and the Barnes (five days) behavioral tasks. After behavioral testing, electrophysiology and tissue collection for metabolomics was performed.

