## Supplementary Data File 2 for "Rescue of hippocampal synaptic plasticity and memory performance by Fingolimod (FTY720) in APP/PS1 model of Alzheimer’s disease is accompanied by correction in metabolism of sphingolipids, polyamines, and phospholipid saturation composition"

| Name | Long name | Formula |
|---|---|---|
| **TMAO synthesis** | Trimethylamine N-oxide synthesis | TMAO / (Betaine + C0 + Choline) |
| **TMAO synthesis (direct)** | Trimethylamine N-oxide synthesis (direct) | TMAO / Choline |
| **Asn synthesis** | Asparagine synthesis | Asn / Asp |
| **Cys synthesis** | Cysteine synthesis | Cys / (Ser + Met) |
| **DLD** | Dihydrolipoamide dehydrogenase deficiency | Pro / Phe |
| **Fischer ratio** | Fischer ratio | (Ile + Leu + Val) / (Phe + Trp + Tyr) |
| **GABR** | Global arginine bioavailability ratio | Arg / (Orn + Cit) |
| **Glutaminase activity** | Glutaminase activity | Glu / Gln |
| **Glutaminolysis rate** | Glutaminolysis rate | (Ala + Asp + Glu + Lac + Suc) / Gln |
| **Gly synthesis** | Glycine synthesis | Gly / Ser |
| **GSH constituents** | Glutathione constituents | Glu + Gly + Cys |
| **MTHFR deficiency** | Methylene tetrahydrofolate reductase deficiency | Met / Phe |
| **PKU** | Phenylketonuria | Tyr / Phe |
| **Ratio of non-essential to essential AAs** | Ratio of non-essential to essential amino acids | (Ala + Arg + Asn + Asp + Cys + Gln + Glu + Gly + Pro + Ser + Tyr) / (His + Ile + Leu + Lys + Met + Phe + Thr + Trp + Val) |
| **Ratio of Pro to Cit** | Ratio of proline to citrulline | Pro / Cit |
| **Sum of AAs** | Sum of amino acids | Ala + Arg + Asn + Asp + Cys + Gln + Glu + Gly + His + Ile + Leu + Lys + Met + Phe + Pro + Ser + Thr + Trp + Tyr + Val |
| **Sum of aromatic AAs** | Sum of aromatic amino acids | Phe + Trp + Tyr |
| **Sum of BCAAs** | Sum of branched-chain amino acids | Ile + Leu + Val |
| **Sum of essential AAs** | Sum of essential amino acids | His + Ile + Leu + Lys + Met + Phe + Thr + Trp + Val |
| **Sum of non-essential AAs** | Sum of non-essential amino acids | Ala + Arg + Asn + Asp + Cys + Gln + Glu + Gly + Pro + Ser + Tyr |
| **Sum of solely glucogenic AAs** | Sum of solely glucogenic amino acids | Ala + Arg + Asn + Asp + Cys + Gln + Glu + Gly + His + Met + Pro + Ser + Thr + Val |
| **Sum of solely ketogenic AAs** | Sum of solely ketogenic amino acids | Leu + Lys |
| **Sum of sulfur-containing AAs** | Sum of sulfur-containing amino acids | Met + Cys |
| **Valinemia** | Valinemia | Val / Phe |
| **1-Met-His synthesis** | 1-Methylhistidine synthesis | 1-Met-His / His |
| **3-Met-His synthesis** | 3-Methylhistidine synthesis | 3-Met-His / (Anserine + Carnosine) |
| **AABA synthesis** | alpha-Aminobutyric acid synthesis | AABA / Thr |
| **Anserine synthesis** | Anserine synthesis | Anserine / Carnosine |
| **Asymmetrical Arg methylation** | Asymmetrical arginine methylation | ADMA / Arg |
| **BABA synthesis** | beta-Aminobutyric acid synthesis | BABA / Glu |
| **Betaine synthesis** | Betaine synthesis | Betaine / Choline |

| Name | Long name | Formula |
|---|---|---|
| **Carnosine synthesis** | Carnosine synthesis | Carnosine / His |
| **Cit synthesis** | Citrulline synthesis | Cit / Orn |
| **CPS deficiency** | Carbamoyl phosphate synthase deficiency | Cit / Phe |
| **Cystine synthesis** | Cystine synthesis | Cystine / Cys |
| **DOPA synthesis** | Dihydroxyphenylalanine synthesis | DOPA / Tyr |
| **HArg synthesis** | Homoarginine synthesis | HArg / (Arg + Lys) |
| **HCys synthesis** | Homocysteine synthesis | HCys / Met |
| **IDO activity** | Indoleamine 2,3-dioxygenase activity | Kynurenine / Trp |
| **Met oxidation** | Methionine oxidation | Met-SO / Met |
| **Muscle protein degradation** | Muscle protein degradation | 1-Met-His / Creatinine |
| **NOS activity** | Nitric oxide synthase activity | Cit / Arg |
| **Orn synthesis** | Ornithine synthesis | Orn / Arg |
| **OTC deficiency** | Ornithine transcarbamylase deficiency | Orn / Cit |
| **Pro hydroxylation** | Proline hydroxylation | (c4-OH-Pro + t4-OH-Pro) / Pro |
| **Ratio of HArg to ADMA** | Ratio of homoarginine to asymmetric dimethylarginine | HArg / ADMA |
| **Ratio of HArg to SDMA** | Ratio of homoarginine to symmetric dimethylarginine | HArg / SDMA |
| **Ratio of SG to hexose** | Ratio of serine and glycine to hexose | (Ser + Gly) / Hexose |
| **Ratio of SGA to hexose** | Ratio of serine, glycine, and alanine to hexose | (Ser + Gly + Ala) / Hexose |
| **Sarcosine synthesis from Gly** | Sarcosine synthesis from glycine | Sarcosine / Gly |
| **Sarcosine synthesis from choline** | Sarcosine synthesis from choline | Sarcosine / Choline |
| **Sum of aminobutyric acids** | Sum of aminobutyric acids | AABA + BABA + GABA |
| **Sum of asym. and sym. Arg methylation** | Sum of asymmetrical and symmetrical arginine methylation | (ADMA + SDMA) / Arg |
| **Sum of betaine and related metabolites** | Sum of betaine and related metabolites | Betaine + PheAlaBetaine + ProBetaine + TrpBetaine |
| **Sum of betaine-related metabolites** | Sum of betaine-related metabolites | PheAlaBetaine + ProBetaine + TrpBetaine |
| **Sum of dimethylated Arg** | Sum of dimethylated arginine | ADMA + SDMA |
| **Symmetrical Arg methylation** | Symmetrical arginine methylation | SDMA / Arg |
| **Taurine synthesis** | Taurine synthesis | Taurine / Cys |
| **Tyr nitration** | Tyrosine nitration | Nitro-Tyr / Tyr |
| **7a-Dehydroxylation of CA** | 7-alpha-Dehydroxylation of cholic acid | DCA / CA |
| **GDCA synthesis from CA** | Glycodeoxycholic acid synthesis from cholic acid | GDCA / CA |
| **GLCA synthesis from CDCA** | Glycolithocholic acid synthesis from chenodeoxycholic acid | GLCA / CDCA |
| **Gly conjugation of CA** | Glycine conjugation of cholic acid | GCA / CA |

| Name | Long name | Formula |
|---|---|---|
| **Gly conjugation of CDCA** | Glycine conjugation of chenodeoxycholic acid | GCDCA / CDCA |
| **Gly conjugation of DCA** | Glycine conjugation of deoxycholic acid | GDCA / DCA |
| **Gly conjugation of primary BAs** | Glycine conjugation of primary bile acids | (GCA + GCDCA) / (CA + CDCA) |
| **Primary BA conjugation** | Ratio of conjugated primary bile acids to unconjugated primary bile acids | (GCA + GCDCA + TCA + TCDCA) / (CA + CDCA) |
| **Ratio of 12a-OH BAs to non-12a-OH BAs** | Ratio of 12-alpha-hydroxylated bile acids to non-12-alpha-hydroxylated bile acids | (CA + DCA + GCA + GDCA + TCA + TDCA) / (CDCA + GCDCA + GLCA + GUDCA + TCDCA + TLCA) |
| **Ratio of CDCA to CA** | Ratio of chenodeoxycholic acid to cholic acid | CDCA / CA |
| **Ratio of primary BAs to BAs** | Ratio of primary bile acids to bile acids | (CA + CDCA + GCA + GCDCA + TCA + TCDCA) / (CA + CDCA + DCA + GCA + GCDCA + GDCA + GLCA + GUDCA + TCA + TCDCA + TDCA + TLCA) |
| **Ratio of secondary BAs to BAs** | Ratio of secondary bile acids to bile acids | (DCA + GDCA + GLCA + GUDCA + TDCA + TLCA) / (CA + CDCA + DCA + GCA + GCDCA + GDCA + GLCA + GUDCA + TCA + TCDCA + TDCA + TLCA) |
| **Secondary BA conjugation** | Ratio of conjugated secondary bile acids to unconjugated secondary bile acids | (GDCA + GLCA + GUDCA + TDCA + TLCA) / DCA |
| **Secondary BA synthesis** | Secondary bile acid synthesis | (DCA + GDCA + GLCA + GUDCA + TDCA + TLCA) / (CA + CDCA + GCA + GCDCA + TCA + TCDCA) |
| **Sum of 12a-OH BAs** | Sum of 12-alpha-hydroxylated bile acids | CA + DCA + GCA + GDCA + TCA + TDCA |
| **Sum of BAs** | Sum of bile acids | CA + CDCA + DCA + GCA + GCDCA + GDCA + GLCA + GUDCA + TCA + TCDCA + TDCA + TLCA |
| **Sum of conjugated BAs** | Sum of conjugated bile acids | GCA + GCDCA + GDCA + GLCA + GUDCA + TCA + TCDCA + TDCA + TLCA |
| **Sum of conjugated primary BAs** | Sum of conjugated primary bile acids | GCA + GCDCA + TCA + TCDCA |
| **Sum of conjugated secondary BAs** | Sum of conjugated secondary bile acids | GDCA + GLCA + GUDCA + TDCA + TLCA |
| **Sum of Gly-conjugated BAs** | Sum of glycine-conjugated bile acids | GCA + GCDCA + GDCA + GLCA + GUDCA |
| **Sum of non-12a-OH BAs** | Sum of non-12-alpha-hydroxylated bile acids | CDCA + GCDCA + GLCA + GUDCA + TCDCA + TLCA |
| **Sum of primary BAs** | Sum of primary bile acids | CA + CDCA + GCA + GCDCA + TCA + TCDCA |
| **Sum of secondary BAs** | Sum of secondary bile acids | DCA + GDCA + GLCA + GUDCA + TDCA + TLCA |
| **Sum of taurine-conjugated BAs** | Sum of taurine-conjugated bile acids | TCA + TCDCA + TDCA + TLCA |
| **Sum of unconjugated BAs** | Sum of unconjugated bile acids | CA + CDCA + DCA |
| **Sum of unconjugated primary BAs** | Sum of unconjugated primary bile acids | CA + CDCA |
| **Taurine conjugation of CA** | Taurine conjugation of cholic acid | TCA / CA |
| **Taurine conjugation of CDCA** | Taurine conjugation of chenodeoxycholic acid | TCDCA / CDCA |
| **Taurine conjugation of DCA** | Taurine conjugation of deoxycholic acid | TDCA / DCA |

| Name | Long name | Formula |
|---|---|---|
| **Taurine conjugation of primary BAs** | Taurine conjugation of primary bile acids | (TCA + TCDCA) / (CA + CDCA) |
| **TDCA synthesis from CA** | Taurodeoxycholic acid synthesis from cholic acid | TDCA / CA |
| **TLCA synthesis from CDCA** | Taurolithocholic acid synthesis from chenodeoxycholic acid | TLCA / CDCA |
| **β-Ala synthesis** | beta-Alanine synthesis | beta-Ala / Carnosine |
| **GABA synthesis** | gamma-Aminobutyric acid synthesis | GABA / Glu |
| **Histamine synthesis** | Histamine synthesis | Histamine / His |
| **PEA synthesis** | Phenylethylamine synthesis | PEA / Phe |
| **Polyamine synthesis** | Polyamine synthesis | (Putrescine + Spermidine + Spermine) / Orn |
| **Putrescine synthesis** | Putrescine synthesis | Putrescine / Orn |
| **Serotonin synthesis** | Serotonin synthesis | Serotonin / Trp |
| **Spermidine synthesis** | Spermidine synthesis | Spermidine / Putrescine |
| **Spermine synthesis** | Spermine synthesis | Spermine / Spermidine |
| **Sum of neurotransmitters** | Sum of neurotransmitters | Dopamine + Histamine + Serotonin |
| **Sum of polyamines** | Sum of polyamines | Putrescine + Spermidine + Spermine |
| **HipAcid synthesis** | Hippuric acid synthesis | HipAcid / Gly |
| **LDH activity** | Lactate dehydrogenase activity | Lac / Hexose |
| **Sum of carboxylic acids** | Sum of carboxylic acids | AconAcid + DiCA(12:0) + DiCA(14:0) + HipAcid + Lac + OH-GlutAcid + Suc |
| **p-Cresol-SO4 synthesis** | para-Cresol sulfate synthesis | p-Cresol-SO4 / Tyr |
| **MAGL activity** | Monoacylglycerol lipase activity | FA 20:4 / MG 20:4 |
| **Ratio of DHA to AA** | Ratio of docosahexaenoic acid to arachidonic acid | FA 22:6 / FA 20:4 |
| **Ratio of DHA to EPA** | Ratio of docosahexaenoic acid to eicosapentaenoic acid | FA 22:6 / FA 20:5 |
| **Ratio of EPA to AA** | Ratio of eicosapentaenoic acid to arachidonic acid | FA 20:5 / FA 20:4 |
| **Ratio of MUFAs to FAs** | Ratio of monounsaturated fatty acids to fatty acids | FA xx:1 / FA xx:x |
| **Ratio of PUFAs to FAs** | Ratio of polyunsaturated fatty acids to fatty acids | FA xx:2-6 / FA xx:x |
| **Ratio of SFAs to FAs** | Ratio of saturated fatty acids to fatty acids | FA xx:0 / FA xx:x |
| **SCD-1 index** | Stearoyl-CoA desaturase-1 index | FA 18:1 / FA 18:0 |
| **Sum of FAs** | Sum of fatty acids | Sum of FA xx:x |
| **Sum of measured ω-3 FAs** | Sum of measured omega-3 fatty acids | FA 20:5 + FA 22:6 |
| **Sum of MUFAs** | Sum of monounsaturated fatty acids | Sum of FA xx:1 |
| **Sum of PUFAs** | Sum of polyunsaturated fatty acids | Sum of FA xx:2-6 |
| **Sum of SFAs** | Sum of saturated fatty acids | Sum of FA xx:0 |
| **Cortisone synthesis** | Cortisone synthesis | Cortisone / Cortisol |

| Name | Long name | Formula |
|---|---|---|
| **Sum of steroid hormones** | Sum of steroid hormones | Cortisol + Cortisone + DHEAS |
| **3-IAA synthesis** | 3-Indoleacetic acid synthesis | 3-IAA / Indole |
| **3-IPA synthesis** | 3-Indolepropionic acid synthesis | 3-IPA / Indole |
| **Indole sulfonation** | Indole sulfonation | Ind-SO4 / Indole |
| **Indole synthesis** | Indole synthesis | Indole / Trp |
| **Sum of indoles** | Sum of indole metabolites | 3-IAA + 3-IPA + Ind-SO4 + Indole |
| **Sum of purines** | Sum of purine derivatives | Hypoxanthine + Xanthine |
| **Xanthine synthesis** | Xanthine synthesis | Xanthine / Hypoxanthine |
| **Sum of choline lipids** | Sum of choline and choline-based lipids | Choline + LPC xx:x + PC (O-)xx:x |
| **2M3HBA** | 2-Methyl-3-hydroxybutyric aciduria | C5-OH (C3-DC-M) / C8 |
| **2MBG** | 2-Methylbutyryl glycinuria | C5 / C3 |
| **3MGA** | 3-Methyl-glutaconic aciduria | C5-OH (C3-DC-M) / C0 |
| **BKT deficiency** | beta-Ketothiolase deficiency | C0 / C5-OH (C3-DC-M) |
| **β-Oxidation** | beta-Oxidation | (C2 + C3) / C0 |
| **CACT deficiency** | Carnitine acylcarnitine translocase deficiency | (C16 + C18) / C0 |
| **Carnitine esterification** | Carnitine esterification | (C2-18 + C2-18:x + Cx-DC + Cx:x-DC + Cx-OH + Cx:x-OH + C5-M-DC) / C0 |
| **Carnitine uptake defect** | Carnitine uptake defect | (C0 + C2 + C3 + C16 + C18 + C18:1) / Cit |
| **CPT-1 deficiency** | Carnitine palmitoyltransferase-1 deficiency | C0 / (C16 + C18) |
| **CPT-2 deficiency** | Carnitine palmitoyltransferase-2 deficiency | (C16 + C18:1) / C2 |
| **EMA** | Ethylmalonic encephalopathy | C4 / C8 |
| **HMG-CoA lyase deficiency** | 3-Hydroxy-3-methylglutaryl-coenzyme A lyase deficiency | C8 / C5-OH (C3-DC-M) |
| **IBD deficiency** | Isobutyryl-coenzyme A dehydrogenase deficiency | C4 / C2 |
| **IVA** | Isovaleric acidemia | C5 / C2 |
| **LCHAD deficiency** | Long-chain 3-hydroxyacyl-coenzyme A dehydrogenase deficiency | C16-OH / C16 |
| **MA** | Malonic aciduria | C3 / C2 |
| **MC deficiency** | Multiple carboxylase deficiency | C16 / C3 |
| **MCAD deficiency** | Medium-chain acyl-coenzyme A dehydrogenase deficiency | C8 / C2 |
| **MCKAT deficiency** | Medium-chain ketoacyl-coenzyme A thiolase deficiency | C8 / C10 |
| **MMA** | Methylmalonic acidemia | C3 / C0 |

| Name | Long name | Formula |
|---|---|---|
| **PA** | Propionic acidemia | C3 / C16 |
| **Ratio of acetylcarnitine to carnitine** | Ratio of acetylcarnitine to carnitine | C2 / C0 |
| **Ratio of AC-OHs to ACs** | Ratio of hydroxylated acylcarnitines to acylcarnitines | (Cx-OH + Cx:x-OH) / (C2-18 + C2-18:x + Cx-DC + Cx:x-DC + Cx-OH + Cx:x-OH + C5-M-DC) |
| **Ratio of ACs to FAs** | Ratio of acylcarnitines to fatty acids | (C2-18 + C2-18:x + Cx-DC + Cx:x-DC + Cx-OH + Cx:x-OH + C5-M-DC) / FA xx:x |
| **Ratio of medium-chain to long-chain ACs** | Ratio of medium-chain acylcarnitines to long-chain acylcarnitines | (C6-12 + C6-12:x + C7-DC + C12-DC) / (C14-18 + C14-18:x + C14:1-OH + C14:2-OH + C16-OH + C16:1-OH + C16:2-OH + C18:1-OH) |
| **Ratio of short-chain to long-chain ACs** | Ratio of short-chain acylcarnitines to long-chain acylcarnitines | (C2-5 + C2-5:x + C3-DC (C4-OH) + C3-OH + C5-DC (C6-OH) + C5-M-DC + C5-OH (C3-DC-M) + C5:1-DC) / (C14-18 + C14-18:x + C14:1-OH + C14:2-OH + C16-OH + C16:1-OH + C16:2-OH + C18:1-OH) |
| **Ratio of short-chain to medium-chain ACs** | Ratio of short-chain acylcarnitines to medium-chain acylcarnitines | (C2-5 + C2-5:x + C3-DC (C4-OH) + C3-OH + C5-DC (C6-OH) + C5-M-DC + C5-OH (C3-DC-M) + C5:1-DC) / (C6-12 + C6-12:x + C7-DC + C12-DC) |
| **SBCAD deficiency** | Short/branched-chain acyl-coenzyme A dehydrogenase deficiency | C5 / C0 |
| **SCAD deficiency** | Short-chain acyl-coenzyme A dehydrogenase deficiency | C4 / C3 |
| **Sum of ACs** | Sum of acylcarnitines | C2-18 + C2-18:x + Cx-DC + Cx:x-DC + Cx-OH + Cx:x-OH + C5-M-DC |
| **Sum of long-chain ACs** | Sum of long-chain acylcarnitines | C14-18 + C14-18:x + C14:1-OH + C14:2-OH + C16-OH + C16:1-OH + C16:2-OH + C18:1-OH |
| **Sum of medium-chain ACs** | Sum of medium-chain acylcarnitines | C6-12 + C6-12:x + C7-DC + C12-DC |
| **Sum of MUFA-ACs** | Sum of monounsaturated fatty acid acylcarnitines | C5:1-DC + Cx:1 + Cx:1-OH |
| **Sum of PUFA-ACs** | Sum of polyunsaturated fatty acid acylcarnitines | Cx:2 + Cx:2-OH |
| **Sum of SFA-ACs** | Sum of saturated fatty acid acylcarnitines | C2-18 + C5-M-DC + Cx-DC + Cx-OH |
| **Sum of short-chain ACs** | Sum of short-chain acylcarnitines | C2-5 + C2-5:x + C3-DC (C4-OH) + C3-OH + C5-DC (C6-OH) + C5-M-DC + C5-OH (C3-DC-M) + C5:1-DC |
| **TFP deficiency** | Mitochondrial tri-functional protein deficiency | C16 / C16-OH |
| **VLCAD deficiency** | Very long-chain acyl-coenzyme A dehydrogenase deficiency | C14:1 / C16 |
| **ω-Oxidation** | omega-Oxidation | (Cx-DC + Cx:x-DC) / (C2-18 + C2-18:x + C5-M-DC + Cx-DC + Cx:x-DC + Cx-OH + Cx:x-OH) |

| Name | Long name | Formula |
|---|---|---|
| **ATX activity** | Autotaxin activity | (LPA xx:x + Choline) / LPC xx:x |
| **MGK activity** | Monoacylglycerol kinase activity | LPA xx:x / MG xx:x |
| **PLA activity (1)** | Phospholipase A activity (1) | LPA xx:x / PA xx:x_xx:x |
| **Ratio of MUFA-LPAs to SFA-LPAs** | Ratio of monounsaturated fatty acid lysophosphatidic acids to saturated fatty acid lysophosphatidic acids | LPA xx:1 / LPA xx:0 |
| **Ratio of PUFA-LPAs to MUFA-LPAs** | Ratio of polyunsaturated fatty acid lysophosphatidic acids to monounsaturated fatty acid lysophosphatidic acids | LPA xx:2-4 / LPA xx:1 |
| **Ratio of PUFA-LPAs to SFA-LPAs** | Ratio of polyunsaturated fatty acid lysophosphatidic acids to saturated fatty acid lysophosphatidic acids | LPA xx:2-4 / LPA xx:0 |
| **Ratio of UFA-LPAs to SFA-LPAs** | Ratio of unsaturated fatty acid lysophosphatidic acids to saturated fatty acid lysophosphatidic acids | LPA xx:1-4 / LPA xx:0 |
| **Sum of LCFA-LPAs** | Sum of long-chain fatty acid lysophosphatidic acids | Sum of LPA 14-18:x |
| **Sum of LPAs** | Sum of lysophosphatidic acids | Sum of LPA xx:x |
| **Sum of MUFA-LPAs** | Sum of monounsaturated fatty acid lysophosphatidic acids | Sum of LPA xx:1 |
| **Sum of PUFA-LPAs** | Sum of polyunsaturated fatty acid lysophosphatidic acids | Sum of LPA xx:2-4 |
| **Sum of SFA-LPAs** | Sum of saturated fatty acid lysophosphatidic acids | Sum of LPA xx:0 |
| **Sum of UFA-LPAs** | Sum of unsaturated fatty acid lysophosphatidic acids | Sum of LPA xx:1-4 |
| **Sum of VLCFA-LPAs** | Sum of very long-chain fatty acid lysophosphatidic acids | Sum of LPA 22:x |
| **DGK activity** | Diacylglycerol kinase activity | PA xx:x_xx:x / DG (O-)xx:x_xx:x |
| **PLD activity** | Phospholipase D activity | (PA xx:x_xx:x + Choline) / PC (O-)xx:x |
| **Ratio of PUFA-PAs to MUFA-PAs** | Ratio of polyunsaturated fatty acid phosphatidic acids to monounsaturated fatty acid phosphatidic acids | (PA xx:2-3_xx:x + PA xx:x_xx:2-4) / (PA xx:1_xx:0-1 + PA xx:0-1_xx:1) |
| **Sum of (L)PAs** | Sum of phosphatidic acids and lysophosphatidic acids | PA xx:x_xx:x + LPA xx:x |
| **Sum of MUFA-PAs** | Sum of monounsaturated fatty acid phosphatidic acids | PA xx:1_xx:0-1 + PA xx:0-1_xx:1 |
| **Sum of PAs** | Sum of phosphatidic acids | Sum of PA xx:x_xx:x |
| **Sum of PUFA-PAs** | Sum of polyunsaturated fatty acid phosphatidic acids | PA xx:2-3_xx:x + PA xx:x_xx:2-4 |
| **PLA2 activity (1)** | Phospholipase A2 activity (1) | LPC xx:x / PC (O-)xx:x |
| **PLA2 activity (2)** | Phospholipase A2 activity (2) | (LPC 16:0 + FA 20:4) / PC 36:4 |

| Name | Long name | Formula |
|---|---|---|
| **PLA2 activity (3)** | Phospholipase A2 activity (3) | (LPC 16:1 + FA 20:4) / PC 36:5 |
| **PLA2 activity (4)** | Phospholipase A2 activity (4) | (LPC 18:0 + FA 20:4) / PC 38:4 |
| **PLA2 activity (5)** | Phospholipase A2 activity (5) | (LPC 18:1 + FA 20:4) / PC 38:5 |
| **PLA2 activity (6)** | Phospholipase A2 activity (6) | (LPC 18:2 + FA 20:4) / PC 38:6 |
| **Ratio of MUFA-LPCs to SFA-LPCs** | Ratio of monounsaturated fatty acid lysophosphatidylcholines to saturated fatty acid lysophosphatidylcholines | LPC xx:1 / LPC xx:0 |
| **Ratio of PUFA-LPCs to MUFA-LPCs** | Ratio of polyunsaturated fatty acid lysophosphatidylcholines to monounsaturated fatty acid lysophosphatidylcholines | LPC xx:2-4 / LPC xx:1 |
| **Ratio of PUFA-LPCs to SFA-LPCs** | Ratio of polyunsaturated fatty acid lysophosphatidylcholines to saturated fatty acid lysophosphatidylcholines | LPC xx:2-4 / LPC xx:0 |
| **Ratio of UFA-LPCs to SFA-LPCs** | Ratio of unsaturated fatty acid lysophosphatidylcholines to saturated fatty acid lysophosphatidylcholines | LPC xx:1-4 / LPC xx:0 |
| **Sum of LCFA-LPCs** | Sum of long-chain fatty acid lysophosphatidylcholines | Sum of LPC 14-20:x |
| **Sum of LPCs** | Sum of lysophosphatidylcholines | Sum of LPC xx:x |
| **Sum of MUFA-LPCs** | Sum of monounsaturated fatty acid lysophosphatidylcholines | Sum of LPC xx:1 |
| **Sum of PUFA-LPCs** | Sum of polyunsaturated fatty acid lysophosphatidylcholines | Sum of LPC xx:2-4 |
| **Sum of SFA-LPCs** | Sum of saturated fatty acid lysophosphatidylcholines | Sum of LPC xx:0 |
| **Sum of UFA-LPCs** | Sum of unsaturated fatty acid lysophosphatidylcholines | Sum of LPC xx:1-4 |
| **Sum of VLCFA-LPCs** | Sum of very long-chain fatty acid lysophosphatidylcholines | Sum of LPC 24-26:x |
| **Ratio of MUFA-PCs to SFA-PCs** | Ratio of monounsaturated fatty acid diacyl-phosphatidylcholines to saturated fatty acid diacyl-phosphatidylcholines | PC xx:1 / PC xx:0 |
| **Ratio of MUFA-PCs O to SFA-PCs O** | Ratio of monounsaturated fatty acid alkylacyl-phosphatidylcholines to saturated fatty acid alkylacyl-phosphatidylcholines | PC O-xx:1 / PC O-xx:0 |

| Name | Long name | Formula |
|---|---|---|
| **Ratio of MUFA-PC (O)s to SFA-PC (O)s** | Ratio of monounsaturated fatty acid phosphatidylcholines to saturated fatty acid phosphatidylcholines | PC (O-)xx:1 / PC (O-)xx:0 |
| **Ratio of PCs to choline** | Ratio of diacyl-phosphatidylcholines to choline | PC xx:x / Choline |
| **Ratio of PCs O to choline** | Ratio of alkylacyl-phosphatidylcholines to choline | PC O-xx:x / Choline |
| **Ratio of PCs O to PCs** | Ratio of alkylacyl-phosphatidylcholines to diacyl-phosphatidylcholines | PC O-xx:x / PC xx:x |
| **Ratio of PC (O)s to choline** | Ratio of phosphatidylcholines to choline | PC (O-)xx:x / Choline |
| **Ratio of PC (O)s to PEs** | Ratio of phosphatidylcholines to diacyl-phosphatidylethanolamines | PC (O-)xx:x / PE xx:x(x) |
| **Ratio of PUFA-PCs to MUFA-PCs** | Ratio of polyunsaturated fatty acid diacyl-phosphatidylcholines to monounsaturated fatty acid diacyl-phosphatidylcholines | PC xx:3-6 / PC xx:1 |
| **Ratio of PUFA-PCs O to MUFA-PCs O** | Ratio of polyunsaturated fatty acid alkylacyl-phosphatidylcholines to monounsaturated fatty acid alkylacyl-phosphatidylcholines | PC O-xx:3-6 / PC O-xx:1 |
| **Ratio of PUFA-PC (O)s to MUFA-PC (O)s** | Ratio of polyunsaturated fatty acid phosphatidylcholines to monounsaturated fatty acid phosphatidylcholines | PC (O-)xx:3-6 / PC (O-)xx:1 |
| **Ratio of PUFA-PCs to SFA-PCs** | Ratio of polyunsaturated fatty acid diacyl-phosphatidylcholines to saturated fatty acid diacyl-phosphatidylcholines | PC xx:3-6 / PC xx:0 |
| **Ratio of PUFA-PCs O to SFA-PCs O** | Ratio of polyunsaturated fatty acid alkylacyl-phosphatidylcholines to saturated fatty acid alkylacyl-phosphatidylcholines | PC O-xx:3-6 / PC O-xx:0 |
| **Ratio of PUFA-PC (O)s to SFA-PC (O)s** | Ratio of polyunsaturated fatty acid phosphatidylcholines to saturated fatty acid phosphatidylcholines | PC (O-)xx:3-6 / PC (O-)xx:0 |
| **Ratio of UFA-PCs to SFA-PCs** | Ratio of unsaturated fatty acid diacyl-phosphatidylcholines to saturated fatty acid diacyl-phosphatidylcholines | PC xx:1-6 / PC xx:0 |
| **Ratio of UFA-PCs O to SFA-PCs O** | Ratio of unsaturated fatty acid alkylacyl-phosphatidylcholines to saturated fatty acid alkylacyl-phosphatidylcholines | PC O-xx:1-6 / PC O-xx:0 |

| Name | Long name | Formula |
|---|---|---|
| **Ratio of UFA-PC (O)s to SFA-PC (O)s** | Ratio of unsaturated fatty acid phosphatidylcholines to saturated fatty acid phosphatidylcholines | PC (O-)xx:1-6 / PC (O-)xx:0 |
| **Sum of (L)PC (O)s** | Sum of phosphatidylcholines and lysophosphatidylcholines | PC (O-)xx:x + LPC xx:x |
| **Sum of MUFA-PCs** | Sum of monounsaturated fatty acid diacyl-phosphatidylcholines | Sum of PC xx:1 |
| **Sum of MUFA-PCs O** | Sum of monounsaturated fatty acid alkylacyl-phosphatidylcholines | Sum of PC O-xx:1 |
| **Sum of MUFA-PC (O)s** | Sum of monounsaturated fatty acid phosphatidylcholines | Sum of PC (O-)xx:1 |
| **Sum of PC (O)s** | Sum of phosphatidylcholines | Sum of PC (O-)xx:x |
| **Sum of PCs** | Sum of diacyl-phosphatidylcholines | Sum of PC xx:x |
| **Sum of PCs O** | Sum of alkylacyl-phosphatidylcholines | Sum of PC O-xx:x |
| **Sum of PUFA-PCs** | Sum of polyunsaturated fatty acid diacyl-phosphatidylcholines | Sum of PC xx:3-6 |
| **Sum of PUFA-PCs O** | Sum of polyunsaturated fatty acid alkylacyl-phosphatidylcholines | Sum of PC O-xx:3-6 |
| **Sum of PUFA-PC (O)s** | Sum of polyunsaturated fatty acid phosphatidylcholines | Sum of PC (O-)xx:3-6 |
| **Sum of SFA-PCs** | Sum of saturated fatty acid diacyl-phosphatidylcholines | Sum of PC xx:0 |
| **Sum of SFA-PCs O** | Sum of saturated fatty acid alkylacyl-phosphatidylcholines | Sum of PC O-xx:0 |
| **Sum of SFA-PC (O)s** | Sum of saturated fatty acid phosphatidylcholines | Sum of PC (O-)xx:0 |
| **Sum of UFA-PCs** | Sum of unsaturated fatty acid diacyl-phosphatidylcholines | Sum of PC xx:1-6 |
| **Sum of UFA-PCs O** | Sum of unsaturated fatty acid alkylacyl-phosphatidylcholines | Sum of PC O-xx:1-6 |
| **Sum of UFA-PC (O)s** | Sum of unsaturated fatty acid phosphatidylcholines | Sum of PC (O-)xx:1-6 |
| **PLA2 activity (7)** | Phospholipase A2 activity (7) | LPE xx:x / PE xx:x(x) |
| **PLA2 activity (8)** | Phospholipase A2 activity (8) | LPE P-xx:x / PE P-xx:x/xx:x |
| **PLA2 activity (9)** | Phospholipase A2 activity (9) | LPE (P-)xx:x / PE (P-)xx:x(x)(/xx:x) |
| **Ratio of LPEs to LPEs P** | Ratio of acyl-lysophosphatidylethanolamines to lysophosphatidylethanolamine plasmalogens | LPE xx:x / LPE P-xx:x |

| Name | Long name | Formula |
|---|---|---|
| **Ratio of MUFA-LPEs to SFA-LPEs** | Ratio of monounsaturated fatty acid acyl-lysophosphatidylethanolamines to saturated fatty acid acyl-lysophosphatidylethanolamines | LPE xx:1 / LPE xx:0 |
| **Ratio of MUFA-LPEs P to SFA-LPEs P** | Ratio of monounsaturated fatty acid lysophosphatidylethanolamine plasmalogens to saturated fatty acid lysophosphatidylethanolamine plasmalogens | LPE P-xx:1 / LPE P-xx:0 |
| **Ratio of MUFA-LPE (P)s to SFA-LPE (P)s** | Ratio of monounsaturated fatty acid lysophosphatidylethanolamines to saturated fatty acid lysophosphatidylethanolamines | LPE (P-)xx:1 / LPE (P-)xx:0 |
| **Ratio of PUFA-LPEs to MUFA-LPEs** | Ratio of polyunsaturated fatty acid acyl-lysophosphatidylethanolamines to monounsaturated fatty acid acyl-lysophosphatidylethanolamines | LPE xx:2-6 / LPE xx:1 |
| **Ratio of PUFA-LPEs P to MUFA-LPEs P** | Ratio of polyunsaturated fatty acid lysophosphatidylethanolamine plasmalogens to monounsaturated fatty acid lysophosphatidylethanolamine plasmalogens | LPE P-xx:2-6 / LPE P-xx:1 |
| **Ratio of PUFA-LPE (P)s to MUFA-LPE (P)s** | Ratio of polyunsaturated fatty acid lysophosphatidylethanolamines to monounsaturated fatty acid lysophosphatidylethanolamines | LPE (P-)xx:2-6 / LPE (P-)xx:1 |
| **Ratio of PUFA-LPEs to SFA-LPEs** | Ratio of polyunsaturated fatty acid acyl-lysophosphatidylethanolamines to saturated fatty acid acyl-lysophosphatidylethanolamines | LPE xx:2-6 / LPE xx:0 |
| **Ratio of PUFA-LPEs P to SFA-LPEs P** | Ratio of polyunsaturated fatty acid lysophosphatidylethanolamine plasmalogens to saturated fatty acid lysophosphatidylethanolamine plasmalogens | LPE P-xx:2-6 / LPE P-xx:0 |
| **Ratio of PUFA-LPE (P)s to SFA-LPE (P)s** | Ratio of polyunsaturated fatty acid lysophosphatidylethanolamines to saturated fatty acid lysophosphatidylethanolamines | LPE (P-)xx:2-6 / LPE (P-)xx:0 |
| **Ratio of UFA-LPEs to SFA-LPEs** | Ratio of unsaturated fatty acid acyl-lysophosphatidylethanolamines to saturated fatty acid acyl-lysophosphatidylethanolamines | LPE xx:1-6 / LPE xx:0 |

| Name | Long name | Formula |
|---|---|---|
| **Ratio of UFA-LPEs P to SFA-LPEs P** | Ratio of unsaturated fatty acid lysophosphatidylethanolamine plasmalogens to saturated fatty acid lysophosphatidylethanolamine plasmalogens | LPE P-xx:1-6 / LPE P-xx:0 |
| **Ratio of UFA-LPE (P)s to SFA-LPE (P)s** | Ratio of unsaturated fatty acid lysophosphatidylethanolamines to saturated fatty acid lysophosphatidylethanolamines | LPE (P-)xx:1-6 / LPE (P-)xx:0 |
| **Sum of LCFA-LPEs** | Sum of long-chain fatty acid acyl-lysophosphatidylethanolamines | Sum of LPE 14-20:x |
| **Sum of LCFA-LPEs P** | Sum of long-chain fatty acid lysophosphatidylethanolamine plasmalogens | Sum of LPE P-14-20:x |
| **Sum of LCFA-LPE (P)s** | Sum of long-chain fatty acid lysophosphatidylethanolamines | Sum of LPE (P-)14-20:x |
| **Sum of LPEs** | Sum of acyl-lysophosphatidylethanolamines | Sum of LPE xx:x |
| **Sum of LPEs P** | Sum of lysophosphatidylethanolamine plasmalogens | Sum of LPE P-xx:x |
| **Sum of LPE (P)s** | Sum of lysophosphatidylethanolamines | Sum of LPE (P-)xx:x |
| **Sum of MUFA-LPEs** | Sum of monounsaturated fatty acid acyl-lysophosphatidylethanolamines | Sum of LPE xx:1 |
| **Sum of MUFA-LPEs P** | Sum of monounsaturated fatty acid lysophosphatidylethanolamine plasmalogens | Sum of LPE P-xx:1 |
| **Sum of MUFA-LPE (P)s** | Sum of monounsaturated fatty acid lysophosphatidylethanolamines | Sum of LPE (P-)xx:1 |
| **Sum of PUFA-LPEs** | Sum of polyunsaturated fatty acid acyl-lysophosphatidylethanolamines | Sum of LPE xx:2-6 |
| **Sum of PUFA-LPEs P** | Sum of polyunsaturated fatty acid lysophosphatidylethanolamine plasmalogens | Sum of LPE P-xx:2-6 |
| **Sum of PUFA-LPE (P)s** | Sum of polyunsaturated fatty acid lysophosphatidylethanolamines | Sum of LPE (P-)xx:2-6 |
| **Sum of SFA-LPEs** | Sum of saturated fatty acid acyl-lysophosphatidylethanolamines | Sum of LPE xx:0 |
| **Sum of SFA-LPEs P** | Sum of saturated fatty acid lysophosphatidylethanolamine plasmalogens | Sum of LPE P-xx:0 |
| **Sum of SFA-LPE (P)s** | Sum of saturated fatty acid lysophosphatidylethanolamines | Sum of LPE (P-)xx:0 |

| Name | Long name | Formula |
|---|---|---|
| **Sum of UFA-LPEs** | Sum of unsaturated fatty acid acyl-lysophosphatidylethanolamines | Sum of LPE xx:1-6 |
| **Sum of UFA-LPEs P** | Sum of unsaturated fatty acid lysophosphatidylethanolamine plasmalogens | Sum of LPE P-xx:1-6 |
| **Sum of UFA-LPE (P)s** | Sum of unsaturated fatty acid lysophosphatidylethanolamines | Sum of LPE (P-)xx:1-6 |
| **Sum of VLCFA-LPEs** | Sum of very long-chain fatty acid acyl-lysophosphatidylethanolamines | Sum of LPE 22-24:x |
| **Sum of VLCFA-LPEs P** | Sum of very long-chain fatty acid lysophosphatidylethanolamine plasmalogens | Sum of LPE P-22:x |
| **Sum of VLCFA-LPE (P)s** | Sum of very long-chain fatty acid lysophosphatidylethanolamines | Sum of LPE (P-)22-24:x |
| **PSD activity** | Phosphatidylserine decarboxylase activity | PE (P-)xx:x(x)(/xx:x) / PS xx:x |
| **Ratio of MUFA-PEs to SFA-PEs** | Ratio of monounsaturated fatty acid diacyl-phosphatidylethanolamines to saturated fatty acid diacyl-phosphatidylethanolamines | PE xx:1 / PE xx:0 |
| **Ratio of MUFA-PEs P to SFA-PEs P** | Ratio of monounsaturated fatty acid phosphatidylethanolamine plasmalogens to saturated fatty acid phosphatidylethanolamine plasmalogens | (PE P-xx:1/xx:0-1 + PE P-xx:0-1/xx:1) / PE P-xx:0/xx:0 |
| **Ratio of MUFA-PE (P)s to SFA-PE (P)s** | Ratio of monounsaturated fatty acid phosphatidylethanolamines to saturated fatty acid phosphatidylethanolamines | (PE xx:1 + PE P-xx:1/xx:0-1 + PE P-xx:0-1/xx:1) / (PE xx:0 + PE P-xx:0/xx:0) |
| **Ratio of PEs to PEs P** | Ratio of diacyl-phosphatidylethanolamines to phosphatidylethanolamine plasmalogens | PE xx:x(x) / PE P-xx:x/xx:x |
| **Ratio of PUFA-PEs to MUFA-PEs** | Ratio of polyunsaturated fatty acid diacyl-phosphatidylethanolamines to monounsaturated fatty acid diacyl-phosphatidylethanolamines | PE xx:3-12 / PE xx:1 |
| **Ratio of PUFA-PEs P to MUFA-PEs P** | Ratio of polyunsaturated fatty acid phosphatidylethanolamine plasmalogens to monounsaturated fatty acid phosphatidylethanolamine plasmalogens | PE P-xx:x/xx:2-6 / (PE P-xx:1/xx:0-1 + PE P-xx:0-1/xx:1) |
| **Ratio of PUFA-PE (P)s to MUFA-PE (P)s** | Ratio of polyunsaturated fatty acid phosphatidylethanolamines to monounsaturated fatty acid phosphatidylethanolamines | (PE xx:3-12 + PE P-xx:x/xx:2-6) / (PE xx:1 + PE P-xx:1/xx:0-1 + PE P-xx:0-1/xx:1) |

| Name | Long name | Formula |
|---|---|---|
| **Ratio of PUFA-PEs to SFA-PEs** | Ratio of polyunsaturated fatty acid diacyl-phosphatidylethanolamines to saturated fatty acid diacyl-phosphatidylethanolamines | PE xx:3-12 / PE xx:0 |
| **Ratio of PUFA-PEs P to SFA-PEs P** | Ratio of polyunsaturated fatty acid phosphatidylethanolamine plasmalogens to saturated fatty acid phosphatidylethanolamine plasmalogens | PE P-xx:x/xx:2-6 / PE P-xx:0/xx:0 |
| **Ratio of PUFA-PE (P)s to SFA-PE (P)s** | Ratio of polyunsaturated fatty acid phosphatidylethanolamines to saturated fatty acid phosphatidylethanolamines | (PE xx:3-12 + PE P-xx:x/xx:2-6) / (PE xx:0 + PE P-xx:0/xx:0) |
| **Ratio of UFA-PEs to SFA-PEs** | Ratio of unsaturated fatty acid diacyl-phosphatidylethanolamines to saturated fatty acid diacyl-phosphatidylethanolamines | PE xx:1-12 / PE xx:0 |
| **Ratio of UFA-PEs P to SFA-PEs P** | Ratio of unsaturated fatty acid phosphatidylethanolamine plasmalogens to saturated fatty acid phosphatidylethanolamine plasmalogens | (PE P-xx:1-6/xx:x + PE P-xx:x/xx:1-6) / PE P-xx:0/xx:0 |
| **Ratio of UFA-PE (P)s to SFA-PE (P)s** | Ratio of unsaturated fatty acid phosphatidylethanolamines to saturated fatty acid phosphatidylethanolamines | (PE xx:1-12 + PE P-xx:1-6/xx:x + PE P-xx:x/xx:1-6) / (PE xx:0 + PE P-xx:0/xx:0) |
| **Sum of MUFA-PEs** | Sum of monounsaturated fatty acid diacyl-phosphatidylethanolamines | Sum of PE xx:1 |
| **Sum of MUFA-PEs P** | Sum of monounsaturated fatty acid phosphatidylethanolamine plasmalogens | PE P-xx:1/xx:0-1 + PE P-xx:0-1/xx:1 |
| **Sum of MUFA-PE (P)s** | Sum of monounsaturated fatty acid phosphatidylethanolamines | PE xx:1 + PE P-xx:1/xx:0-1 + PE P-xx:0-1/xx:1 |
| **Sum of PEs** | Sum of diacyl-phosphatidylethanolamines | Sum of PE xx:x(x) |
| **Sum of PEs P** | Sum of phosphatidylethanolamine plasmalogens | Sum of PE P-xx:x/xx:x |
| **Sum of PE (P)s** | Sum of phosphatidylethanolamines | PE xx:x(x) + PE P-xx:x/xx:x |
| **Sum of (L)PEs** | Sum of diacyl-phosphatidylethanolamines and acyl-lysophosphatidylethanolamines | Sum of (L)PE xx:x(x) |
| **Sum of (L)PEs P** | Sum of phosphatidylethanolamine plasmalogens and lysophosphatidylethanolamine plasmalogens | PE P-xx:x/xx:x + LPE P-xx:x |
| **Sum of (L)PE (P)s** | Sum of phosphatidylethanolamines and lysophosphatidylethanolamines | PE xx:x(x) + PE P-xx:x/xx:x + LPE xx:x + LPE P-xx:x |

| Name | Long name | Formula |
|---|---|---|
| **Sum of PUFA-PEs** | Sum of polyunsaturated fatty acid diacyl-phosphatidylethanolamines | Sum of PE xx:3-12 |
| **Sum of PUFA-PEs P** | Sum of polyunsaturated fatty acid phosphatidylethanolamine plasmalogens | PE P-xx:x/xx:2-6 |
| **Sum of PUFA-PE (P)s** | Sum of polyunsaturated fatty acid phosphatidylethanolamines | PE xx:3-12 + PE P-xx:x/xx:2-6 |
| **Sum of SFA-PEs** | Sum of saturated fatty acid diacyl-phosphatidylethanolamines | Sum of PE xx:0 |
| **Sum of SFA-PEs P** | Sum of saturated fatty acid phosphatidylethanolamine plasmalogens | PE P-xx:0/xx:0 |
| **Sum of SFA-PE (P)s** | Sum of saturated fatty acid phosphatidylethanolamines | PE xx:0 + PE P-xx:0/xx:0 |
| **Sum of UFA-PEs** | Sum of unsaturated fatty acid diacyl-phosphatidylethanolamines | Sum of PE xx:1-12 |
| **Sum of UFA-PEs P** | Sum of unsaturated fatty acid phosphatidylethanolamine plasmalogens | PE P-xx:1-6/xx:x + PE P-xx:x/xx:1-6 |
| **Sum of UFA-PE (P)s** | Sum of unsaturated fatty acid phosphatidylethanolamines | PE xx:1-12 + PE P-xx:1-6/xx:x + PE P-xx:x/xx:1-6 |
| **PLA2 activity (10)** | Phospholipase A2 activity (10) | LPG xx:x / PG xx:x_xx:x |
| **Ratio of MUFA-LPGs to SFA-LPGs** | Ratio of monounsaturated fatty acid lysophosphatidylglycerols to saturated fatty acid lysophosphatidylglycerols | LPG xx:1 / LPG xx:0 |
| **Ratio of UFA-LPGs to SFA-LPGs** | Ratio of unsaturated fatty acid lysophosphatidylglycerols to saturated fatty acid lysophosphatidylglycerols | LPG xx:1-2 / LPG xx:0 |
| **Sum of LPGs** | Sum of lysophosphatidylglycerols | Sum of LPG xx:x |
| **Sum of MUFA-LPGs** | Sum of monounsaturated fatty acid lysophosphatidylglycerols | Sum of LPG xx:1 |
| **Sum of SFA-LPGs** | Sum of saturated fatty acid lysophosphatidylglycerols | Sum of LPG xx:0 |
| **Sum of UFA-LPGs** | Sum of unsaturated fatty acid lysophosphatidylglycerols | Sum of LPG xx:1-2 |
| **Ratio of MUFA-PGs to SFA-PGs** | Ratio of monounsaturated fatty acid phosphatidylglycerols to saturated fatty acid phosphatidylglycerols | (PG xx:1_xx:0-1 + PG xx:0-1_xx:1) / (PG xx:0_xx:0) |

| Name | Long name | Formula |
|---|---|---|
| **Ratio of PUFA-PGs to MUFA-PGs** | Ratio of polyunsaturated fatty acid phosphatidylglycerols to monounsaturated fatty acid phosphatidylglycerols | (PG xx:2-6_xx:x + PG xx:x_xx:2-6) / (PG xx:1_xx:0-1 + PG xx:0-1_xx:1) |
| **Ratio of PUFA-PGs to SFA-PGs** | Ratio of polyunsaturated fatty acid phosphatidylglycerols to saturated fatty acid phosphatidylglycerols | (PG xx:2-6_xx:x + PG xx:x_xx:2-6) / PG xx:0_xx:0 |
| **Ratio of UFA-PGs to SFA-PGs** | Ratio of unsaturated fatty acid phosphatidylglycerols to saturated fatty acid phosphatidylglycerols | PG xx:1-6_xx:1-6 / PG xx:0_xx:0 |
| **Sum of (L)PGs** | Sum of phosphatidylglycerols and lysophosphatidylglycerols | PG xx:x_xx:x + LPG xx:x |
| **Sum of PGs** | Sum of phosphatidylglycerols | Sum of PG xx:x_xx:x |
| **Sum of MUFA-PGs** | Sum of monounsaturated fatty acid phosphatidylglycerols | PG xx:1_xx:0-1 + PG xx:0-1_xx:1 |
| **Sum of PUFA-PGs** | Sum of polyunsaturated fatty acid phosphatidylglycerol | PG xx:2-6_xx:x + PG xx:x_xx:2-6 |
| **Sum of SFA-PGs** | Sum of saturated fatty acid phosphatidylglycerols | Sum of PG xx:0_xx:0 |
| **Sum of UFA-PGs** | Sum of unsaturated fatty acid phosphatidylglycerols | Sum of PG xx:1-6_xx:1-6 |
| **Sum of VLCFA-LPIs** | Sum of very-long chain fatty acid lysophosphatidylinositols | Sum of LPI 22:x |
| **Sum of UFA-LPIs** | Sum of unsaturated fatty acid lysophosphatidylinositols | Sum of LPI xx:1-4 |
| **Sum of SFA-LPIs** | Sum of saturated fatty acid lysophosphatidylinositols | Sum of LPI xx:0 |
| **Sum of PUFA-LPIs** | Sum of polyunsaturated fatty acid lysophosphatidylinositols | Sum of LPI xx:2-4 |
| **Sum of MUFA-LPIs** | Sum of monounsaturated fatty acid lysophosphatidylinositols | Sum of LPI xx:1 |
| **Sum of LPIs** | Sum of lysophosphatidylinositols | Sum of LPI xx:x |
| **Sum of LCFA-LPIs** | Sum of long-chain fatty acid lysophosphatidylinositols | Sum of LPI 14-20:x |
| **Ratio of UFA-LPIs to SFA-LPIs** | Ratio of unsaturated fatty acid lysophosphatidylinositols to saturated fatty acid lysophosphatidylinositols | LPI xx:1-4 / LPI xx:0 |
| **Ratio of PUFA-LPIs to SFA-LPIs** | Ratio of polyunsaturated fatty acid lysophosphatidylinositols to saturated fatty acid lysophosphatidylinositols | LPI xx:2-4 / LPI xx:0 |

| Name | Long name | Formula |
| --- | --- | --- |
| **Ratio of PUFA-LPIs to MUFA-LPIs** | Ratio of polyunsaturated fatty acid lysophosphatidylinositols to monounsaturated fatty acid lysophosphatidylinositols | LPI xx:2-4 / LPI xx:1 |
| **Ratio of MUFA-LPIs to SFA-LPIs** | Ratio of monounsaturated fatty acid lysophosphatidylinositols to saturated fatty acid lysophosphatidylinositols | LPI xx:1 / LPI xx:0 |
| **PLA activity (2)** | Phospholipase A activity (2) | LPI xx:x / PI xx:x_xx:x |
| **PLA activity (3)** | Phospholipase A activity (3) | (LPI 16:0 + FA 20:4) / PI 16:0_20:4 |
| **PLA activity (4)** | Phospholipase A activity (4) | (LPI 18:0 + FA 20:4) / PI 18:0_20:4 |
| **PLA activity (5)** | Phospholipase A activity (5) | (LPI 18:1 + FA 20:4) / PI 18:1_20:4 |
| **PLA activity (6)** | Phospholipase A activity (6) | (LPI 18:2 + FA 20:4) / PI 18:2_20:4 |
| **LPIAT1 activity (1)** | Lysophosphatidylinositol acyltransferase 1 activity (1) | PI 16:0_20:4 / (LPI 16:0 + FA 20:4) |
| **LPIAT1 activity (2)** | Lysophosphatidylinositol acyltransferase 1 activity (2) | PI 18:0_20:4 / (LPI 18:0 + FA 20:4) |
| **LPIAT1 activity (3)** | Lysophosphatidylinositol acyltransferase 1 activity (3) | PI 18:1_20:4 / (LPI 18:1 + FA 20:4) |
| **LPIAT1 activity (4)** | Lysophosphatidylinositol acyltransferase 1 activity (4) | PI 18:2_20:4 / (LPI 18:2 + FA 20:4) |
| **PI synthesis** | Phosphatidylinositol synthesis | PI xx:x_xx:x / PA xx:x_xx:x |
| **Sum of UFA-PIs** | Sum of unsaturated fatty acid phosphatidylinositols | PI xx:1-2_xx:x + PI xx:x_xx:1-6 |
| **Sum of PUFA-PIs** | Sum of polyunsaturated fatty acid phosphatidylinositols | PI xx:2_xx:x + PI xx:x_xx:2-6 |
| **Sum of (L)PIs** | Sum of phosphatidylinositols and lysophosphatidylinositols | PI xx:x_xx:x + LPI xx:x |
| **Sum of MUFA-PIs** | Sum of monounsaturated fatty acid phosphatidylinositols | PI xx:1_xx:0 + PI xx:0_xx:1 + PI xx:1_xx:1 |
| **Sum of SFA-PIs** | Sum of saturated fatty acid phosphatidylinositols | Sum of PI xx:0_xx:0 |
| **Ratio of UFA-PIs to SFA-PIs** | Ratio of unsaturated fatty acid phosphatidylinositols to saturated fatty acid phosphatidylinositols | (PI xx:1-2_xx:x + PI xx:x_xx:1-6) / PI xx:0_xx:0 |
| **Ratio of PUFA-PIs to SFA-PIs** | Ratio of polyunsaturated fatty acid phosphatidylinositols to saturated fatty acid phosphatidylinositols | (PI xx:2_xx:x + PI xx:x_xx:2-6) / PI xx:0_xx:0 |
| **Ratio of PUFA-PIs to MUFA-PIs** | Ratio of polyunsaturated fatty acid phosphatidylinositols to monounsaturated fatty acid phosphatidylinositols | (PI xx:2_xx:x + PI xx:x_xx:2-6) / (PI xx:1_xx:0 + PI xx:0_xx:1 + PI xx:1_xx:1) |
| **Ratio of MUFA-PIs to SFA-PIs** | Ratio of monounsaturated fatty acid phosphatidylinositols to saturated fatty acid phosphatidylinositols | (PI xx:1_xx:0 + PI xx:0_xx:1 + PI xx:1_xx:1) / PI xx:0_xx:0 |

| Name | Long name | Formula |
|---|---|---|
| **Sum of PIs** | Sum of phosphatidylinositols | Sum of PI xx:x_xx:x |
| **Ratio of PI 18:0_20:4 to PIs** | Ratio of PI 18:0_20:4 to phosphatidylinositols | PI 18:0_20:4 / PI xx:x_xx:x |
| **PLA1A activity** | Phospholipase A1 member A activity | LPS xx:x / PS xx:x |
| **Ratio of MUFA-LPSs to SFA-LPSs** | Ratio of monounsaturated fatty acid lysophosphatidylserines to saturated fatty acid lysophosphatidylserines | LPS xx:1 / LPS xx:0 |
| **Ratio of PUFA-LPSs to MUFA-LPSs** | Ratio of polyunsaturated fatty acid lysophosphatidylserines to monounsaturated fatty acid lysophosphatidylserines | LPS xx:2-6 / LPS xx:1 |
| **Ratio of PUFA-LPSs to SFA-LPSs** | Ratio of polyunsaturated fatty acid lysophosphatidylserines to saturated fatty acid lysophosphatidylserines | LPS xx:2-6 / LPS xx:0 |
| **Ratio of UFA-LPSs to SFA-LPSs** | Ratio of unsaturated fatty acid lysophosphatidylserines to saturated fatty acid lysophosphatidylserines | LPS xx:1-6 / LPS xx:0 |
| **Sum of LCFA-LPSs** | Sum of long-chain fatty acid lysophosphatidylserines | Sum of LPS 16-20:x |
| **Sum of LPSs** | Sum of lysophosphatidylserines | Sum of LPS xx:x |
| **Sum of MUFA-LPSs** | Sum of monounsaturated fatty acid lysophosphatidylserines | Sum of LPS xx:1 |
| **Sum of PUFA-LPSs** | Sum of polyunsaturated fatty acid lysophosphatidylserines | Sum of LPS xx:2-6 |
| **Sum of SFA-LPSs** | Sum of saturated fatty acid lysophosphatidylserines | Sum of LPS xx:0 |
| **Sum of UFA-LPSs** | Sum of unsaturated fatty acid lysophosphatidylserines | Sum of LPS xx:1-6 |
| **Sum of VLCFA-LPSs** | Sum of very long-chain fatty acid lysophosphatidylserines | Sum of LPS 22:x |
| **PSS activity** | Phosphatidylserine synthase activity | PS xx:x / Ser |
| **PSS1 activity** | Phosphatidylserine synthase I activity | (PS xx:x + Choline) / (PC (O-)xx:x + Ser) |
| **PSS2 activity** | Phosphatidylserine synthase II activity | PS xx:x / (PE xx:x(x) + PE P-xx:x/xx:x + Ser) |
| **Ratio of MUFA-PSs to SFA-PSs** | Ratio of monounsaturated fatty acid phosphatidylserines to saturated fatty acid phosphatidylserines | PS xx:1 / PS xx:0 |
| **Ratio of PUFA-PSs to MUFA-PSs** | Ratio of polyunsaturated fatty acid phosphatidylserines to monounsaturated fatty acid phosphatidylserines | PS xx:3-8 / PS xx:1 |

| Name | Long name | Formula |
|---|---|---|
| **Ratio of PUFA-PSs to SFA-PSs** | Ratio of polyunsaturated fatty acid phosphatidylserines to saturated fatty acid phosphatidylserines | PS xx:3-8 / PS xx:0 |
| **Ratio of UFA-PSs to SFA-PSs** | Ratio of unsaturated fatty acid phosphatidylserines to saturated fatty acid phosphatidylserines | PS xx:1-8 / PS xx:0 |
| **Sum of MUFA-PSs** | Sum of monounsaturated fatty acid phosphatidylserines | Sum of PS xx:1 |
| **Sum of PSs** | Sum of phosphatidylserines | Sum of PS xx:x |
| **Sum of (L)PSs** | Sum of phosphatidylserines and lysophosphatidylserines | PS xx:x + LPS xx:x |
| **Sum of PUFA-PSs** | Sum of polyunsaturated fatty acid phosphatidylserines | Sum of PS xx:3-8 |
| **Sum of SFA-PSs** | Sum of saturated fatty acid phosphatidylserines | Sum of PS xx:0 |
| **Sum of UFA-PSs** | Sum of unsaturated fatty acid phosphatidylserines | Sum of PS xx:1-8 |
| **LPP3 activity (1)** | Lipid phosphate phosphatase 3 activity (1) | SPB d14:1 / SPBP d14:1 |
| **LPP3 activity (2)** | Lipid phosphate phosphatase 3 activity (2) | SPB d16:1 / SPBP d16:1 |
| **LPP3 activity (3)** | Lipid phosphate phosphatase 3 activity (3) | SPB d17:1 / SPBP d17:1 |
| **LPP3 activity (4)** | Lipid phosphate phosphatase 3 activity (4) | SPB d18:1 / SPBP d18:1 |
| **LPP3 activity (5)** | Lipid phosphate phosphatase 3 activity (5) | SPB dxx:1 / SPBP dxx:1 |
| **Ratio of Spho to Spha** | Ratio of sphingosines to sphinganines | SPB dxx:1 / SPB dxx:0 |
| **Ratio of SphoPs to SphaPs** | Ratio of sphingosine phosphates to sphinganine phosphates | SPBP dxx:1 / SPBP dxx:0 |
| **Ratio of Spho(P)s to Spha(P)s** | Ratio of sphingosines and their phosphates to sphinganines and their phosphates | SPB(P) dxx:1 / SPB(P) dxx:0 |
| **SphK activity (1)** | Sphingosine kinase activity (1) | SPBP d14:1 / SPB d14:1 |
| **SphK activity (2)** | Sphingosine kinase activity (2) | SPBP d16:1 / SPB d16:1 |
| **SphK activity (3)** | Sphingosine kinase activity (3) | SPBP d17:1 / SPB d17:1 |
| **SphK activity (4)** | Sphingosine kinase activity (4) | SPBP d18:1 / SPB d18:1 |
| **SphK activity (5)** | Sphingosine kinase activity (5) | SPBP dxx:1 / SPB dxx:1 |
| **SphK activity (6)** | Sphingosine kinase activity (6) | SPBP d14:0 / SPB d14:0 |
| **SphK activity (7)** | Sphingosine kinase activity (7) | SPBP d16:0 / SPB d16:0 |
| **SphK activity (8)** | Sphingosine kinase activity (8) | SPBP d17:0 / SPB d17:0 |
| **SphK activity (9)** | Sphingosine kinase activity (9) | SPBP d18:0 / SPB d18:0 |
| **SphK activity (10)** | Sphingosine kinase activity (10) | SPBP dxx:0 / SPB dxx:0 |
| **SphK activity (11)** | Sphingosine kinase activity (11) | SPBP dxx:x / SPB dxx:x |

| Name | Long name | Formula |
|---|---|---|
| **Spho synthesis** | Sphingosine synthesis | SPB dxx:1 / Cer dxx:1/xx:x |
| **Sum of Spha** | Sum of sphinganines | Sum of SPB dxx:0 |
| **Sum of SphaPs** | Sum of sphinganine phosphates | Sum of SPBP dxx:0 |
| **Sum of Spha(P)s** | Sum of sphinganines and their phosphates | Sum of SPB(P) dxx:0 |
| **Sum of SPBs** | Sum of sphingoid bases | Sum of SPB dxx:x |
| **Sum of SPBPs** | Sum of sphingoid base phosphates | Sum of SPBP dxx:x |
| **Sum of SLs** | Sum of sphingolipids | SPB(P) dxx:x + SM xx:x + Cer(P) dxx:x/xx:x(-OH) + Hex[-23]Cer dxx:x/xx:x |
| **Sum of Spho** | Sum of sphingosines | Sum of SPB dxx:1 |
| **Sum of SphoPs** | Sum of sphingosine phosphates | Sum of SPBP dxx:1 |
| **Sum of Spho(P)s** | Sum of sphingosines and their phosphates | Sum of SPB(P) dxx:1 |
| **Sum of SPB(P)s** | Sum of sphingoid bases and their phosphates | Sum of SPB(P) dxx:x |
| **Ratio of OC-FA SMs to EC-FA SMs** | Ratio of odd-chain fatty acid sphingomyelins to even-chain fatty acid sphingomyelins | (SM x1:x + SM x3:x + SM x5:x) / (SM x2:x + SM x4:x + SM x6:x + SM x8:x + SM x0:x) |
| **Ratio of SMs to Cer** | Ratio of sphingomyelins to ceramides | SM xx:x / Cer d1x:x/xx:x(-OH) |
| **Ratio of SMs to PC (O)s** | Ratio of sphingomyelins to phosphatidylcholines | SM xx:x / PC (O-)xx:x |
| **Sum of LCFA-SMs** | Sum of long-chain fatty acid sphingomyelins | SM 33-36:x |
| **Sum of EC-FA SMs** | Sum of even-chain fatty acid sphingomyelins | SM x2:x + SM x4:x + SM x6:x + SM x8:x + SM x0:x |
| **Sum of OC-FA SMs** | Sum of odd-chain fatty acid sphingomyelins | SM x1:x + SM x3:x + SM x5:x |
| **Sum of SMs** | Sum of sphingomyelins | Sum of SM xx:x |
| **Sum of VLCFA-SMs** | Sum of very long-chain fatty acid sphingomyelins | Sum of SM 42-44:x |
| **CERK activity** | Ceramide kinase activity | CerP d18:1/16:0 / Cer d18:1/16:0 |
| **CPP activity** | Ceramide phosphate phosphatase activity | Cer d18:1/16:0 / CerP d18:1/16:0 |
| **SMase activity** | Sphingomyelinase activity | Cer d1x:x/xx:x(-OH) / SM xx:x |
| **Ratio of Cer to DH-Cer** | Ratio of ceramides to dihydroceramides | Cer d1x:x/xx:x(-OH) / Cer d18:0/xx:x(-OH) |
| **Ratio of Cer to Spho** | Ratio of ceramides to sphingosines | Cer d1x:x/xx:x(-OH) / SPB dxx:1 |
| **Ratio of DH-Cer to Spha** | Ratio of dihydroceramides to sphinganines | Cer d18:0/xx:x(-OH) / SPB dxx:0 |
| **Ratio of glycosylcer to Cer** | Ratio of glycosylceramides to ceramides | (Hex-Cer d1x:x/xx:x + Hex2Cer d18:1/xx:x + Hex3Cer d18:1/xx:x) / Cer d1x:x/xx:x(-OH) |
| **Ratio of Hex2Cer to Cer** | Ratio of dihexosylceramides to ceramides | Hex2Cer d18:1/xx:x / Cer d1x:x/xx:x(-OH) |
| **Ratio of Hex2Cer to Hex-Cer** | Ratio of dihexosylceramides to hexosylceramides | Hex2Cer d18:1/xx:x / Hex-Cer d1x:x/xx:x |
| **Ratio of Hex3Cer to Cer** | Ratio of trihexosylceramides to ceramides | Hex3Cer d18:1/xx:x / Cer d1x:x/xx:x(-OH) |
| **Ratio of Hex3Cer to Hex2Cer** | Ratio of trihexosylceramides to dihexosylceramides | Hex3Cer d18:1/xx:x / Hex2Cer d18:1/xx:x |
| **Ratio of Hex3Cer to Hex-Cer** | Ratio of trihexosylceramides to hexosylceramides | Hex3Cer d18:1/xx:x / Hex-Cer d1x:x/xx:x |
| **Ratio of Hex-Cer to Cer** | Ratio of hexosylceramides to ceramides | Hex-Cer d1x:x/xx:x / Cer d1x:x/xx:x(-OH) |

| Name | Long name | Formula |
|---|---|---|
| **Sum of Cer** | Sum of ceramides | Cer d1x:x/xx:x(-OH) |
| **Sum of DH-Cer** | Sum of dihydroceramides | Sum of Cer d18:0/xx:x(-OH) |
| **Sum of glycosylcer** | Sum of glycosylceramides | Hex-Cer d1x:x/xx:x + Hex2Cer d18:1/xx:x + Hex3Cer d18:1/xx:x |
| **Sum of Hex2Cer** | Sum of dihexosylceramides | Sum of Hex2Cer d18:1/xx:x |
| **Sum of Hex3Cer** | Sum of trihexosylceramides | Sum of Hex3Cer d18:1/xx:x |
| **Sum of Hex-Cer** | Sum of hexosylceramides | Sum of Hex-Cer d1x:x/xx:x |
| **Sum of LCFA-Cer** | Sum of long-chain fatty acid ceramides | Cer d1x:1/14-20:x(-OH) |
| **Sum of LCFA-DH-Cer** | Sum of long-chain fatty acid dihydroceramides | Sum of Cer d18:0/18-20:0(-OH) |
| **Sum of LCFA-glycosylcer** | Sum of long-chain fatty acid glycosylceramides | Hex-Cer d18:x/14-20:x + Hex2Cer d18:1/14-20:0 + Hex3Cer d18:1/16-20:0 |
| **Sum of VLCFA-Cer** | Sum of very long-chain fatty acid ceramides | Sum of Cer d1x:x/22-26:x |
| **Sum of VLCFA-DH-Cer** | Sum of very long-chain fatty acid dihydroceramides | Sum of Cer d18:0/22-26:x(-OH) |
| **Sum of VLCFA-glycosylcer** | Sum of very long-chain fatty acid glycosylceramides | Hex-Cer d1x:x/22-26:x + Hex2Cer d18:1/22-26:x + Hex3Cer d18:1/22-26:x |
| **Sum of CEs** | Sum of cholesteryl esters | Sum of CE xx:x |
| **Sum of LCFA-CEs** | Sum of long-chain fatty acid cholesteryl esters | Sum of CE 14-20:x |
| **Sum of MUFA-CEs** | Sum of monounsaturated fatty acid cholesteryl esters | Sum of CE xx:1 |
| **Sum of PUFA-CEs** | Sum of polyunsaturated fatty acid cholesteryl esters | Sum of CE xx:2-6 |
| **Sum of saturated CEs** | Sum of saturated cholesteryl esters | Sum of CE xx:0 |
| **Sum of VLCFA-CEs** | Sum of very long-chain fatty acid cholesteryl esters | Sum of CE 22:x |
| **LPP3 activity (6)** | Lipid phosphate phosphatase 3 activity (6) | MG xx:x / LPA xx:x |
| **Ratio of MGs to DGs** | Ratio of monoglycerides to diglycerides | MG xx:x / DG (O-)xx:x_xx:x |
| **Ratio of MGs to FAs** | Ratio of monoglycerides to fatty acids | MG xx:x / FA xx:x |
| **Ratio of MGs to TGs** | Ratio of monoglycerides to triglycerides | MG xx:x / TG xx:x_xx:x |
| **Sum of GLs** | Sum of glycerolipids | MG xx:x + DG (O-)xx:x_xx:x + TG xx:x_xx:x |
| **Sum of MGs** | Sum of monoglycerides | Sum of MG xx:x |
| **Sum of MUFA-MGs** | Sum of monounsaturated fatty acid monoglycerides | Sum of MG xx:1 |
| **Sum of PUFA-MGs** | Sum of polyunsaturated fatty acid monoglycerides | Sum of MG xx:2-6 |
| **LPP3 activity (7)** | Lipid phosphate phosphatase 3 activity (7) | DG (O-)xx:x_xx:x / PA xx:x_xx:x |
| **PLC activity** | Phospholipase C activity | DG (O-)xx:x_xx:x / PI xx:x_xx:x |
| **Ratio of DGs to FAs** | Ratio of diglycerides to fatty acids | DG (O-)xx:x_xx:x / FA xx:x |
| **Ratio of DGs to TGs** | Ratio of diglycerides to triglycerides | DG (O-)xx:x_xx:x / TG xx:x_xx:x |
| **Sum of DGs** | Sum of diglycerides | Sum of DG (O-)xx:x_xx:x |
| **Sum of MUFA-DGs** | Sum of monounsaturated fatty acid diglycerides | DG (O-)xx:1_xx:0-1 + DG (O-)xx:0-1_xx:1 |

| Name | Long name | Formula |
|---|---|---|
| **Sum of PUFA-DGs** | Sum of polyunsaturated fatty acid diglycerides | DG (O-)xx:2-3_xx:x + DG (O-)xx:x_xx:2-6 |
| **Sum of SFA-DGs** | Sum of saturated fatty acid diglycerides | Sum of DG xx:0_xx:0 |
| **Sum of UFA-DGs** | Sum of unsaturated fatty acid diglycerides | DG (O-)xx:x_xx:1-6 + DG (O-)xx:1-3_xx:x |
| **Ratio of TGs to FAs** | Ratio of triglycerides to fatty acids | TG xx:x_xx:x / FA xx:x |
| **Sum of MUFA-TGs** | Sum of monounsaturated fatty acid triglycerides | TG xx:1_xx:0-1 + TG xx:0-1_xx:1 |
| **Sum of PUFA-TGs** | Sum of polyunsaturated fatty acid triglycerides | TG xx:2-6_xx:x + TG xx:x_xx:3-8 |
| **Sum of SFA-TGs** | Sum of saturated fatty acid triglycerides | Sum of TG xx:0_xx:0 |
| **Sum of TGs** | Sum of triglycerides | Sum of TG xx:x_xx:x |
| **Sum of UFA-TGs** | Sum of unsaturated fatty acid triglycerides | TG xx:x_xx:1-8 + TG xx:1-6_xx:x |
