## Supplementary figures and images for "Rescue of hippocampal synaptic plasticity and memory performance by Fingolimod (FTY720) in APP/PS1 model of Alzheimer’s disease is accompanied by correction in metabolism of sphingolipids, polyamines, and phospholipid saturation composition"

### Supplementary Figure 2

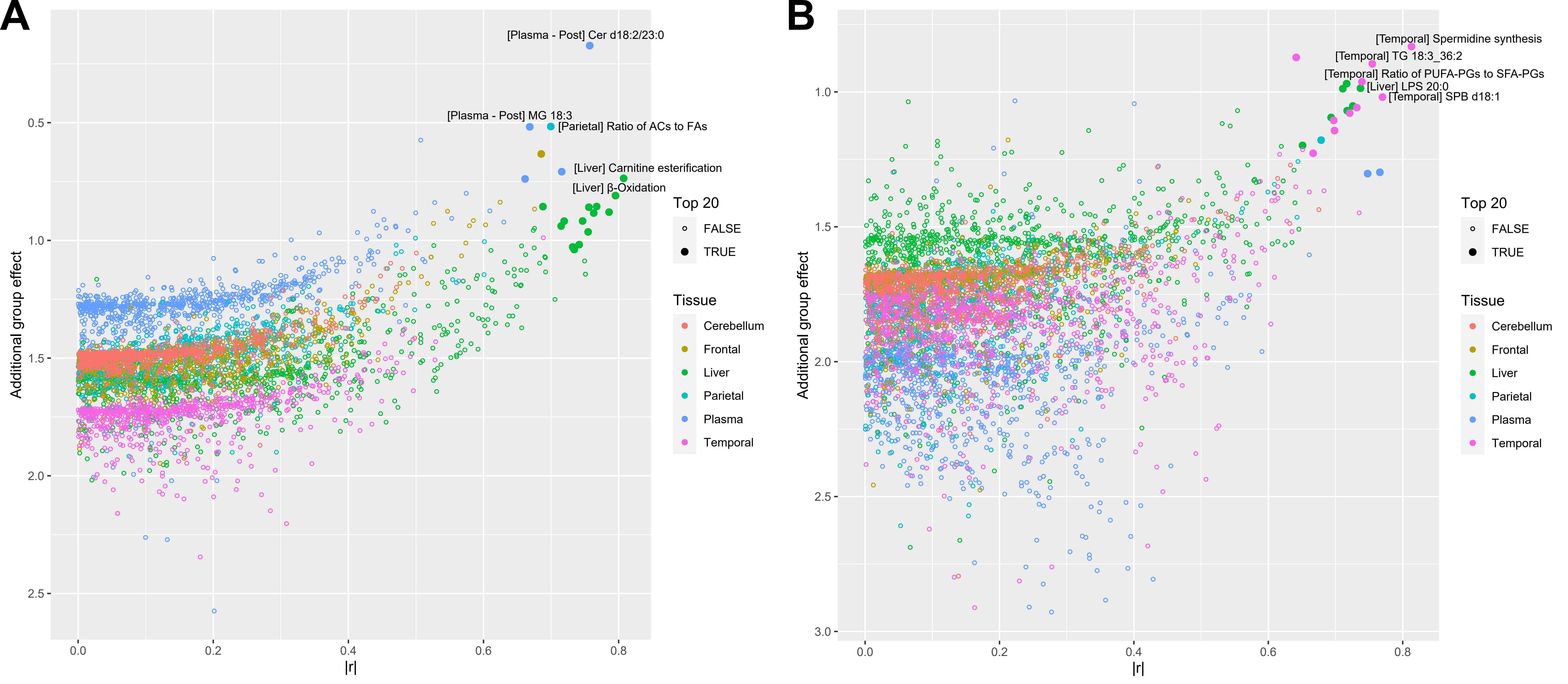

### Supplementary Figure 3

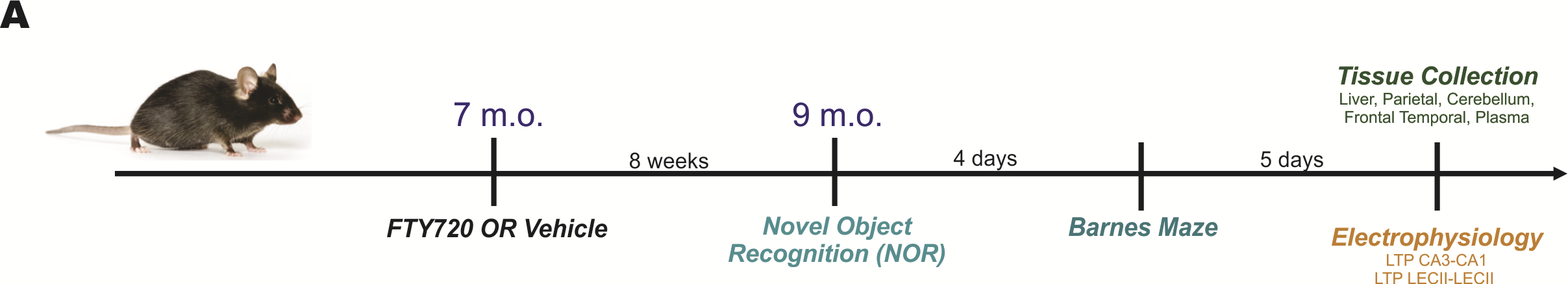

### Suuplementary Figure 1

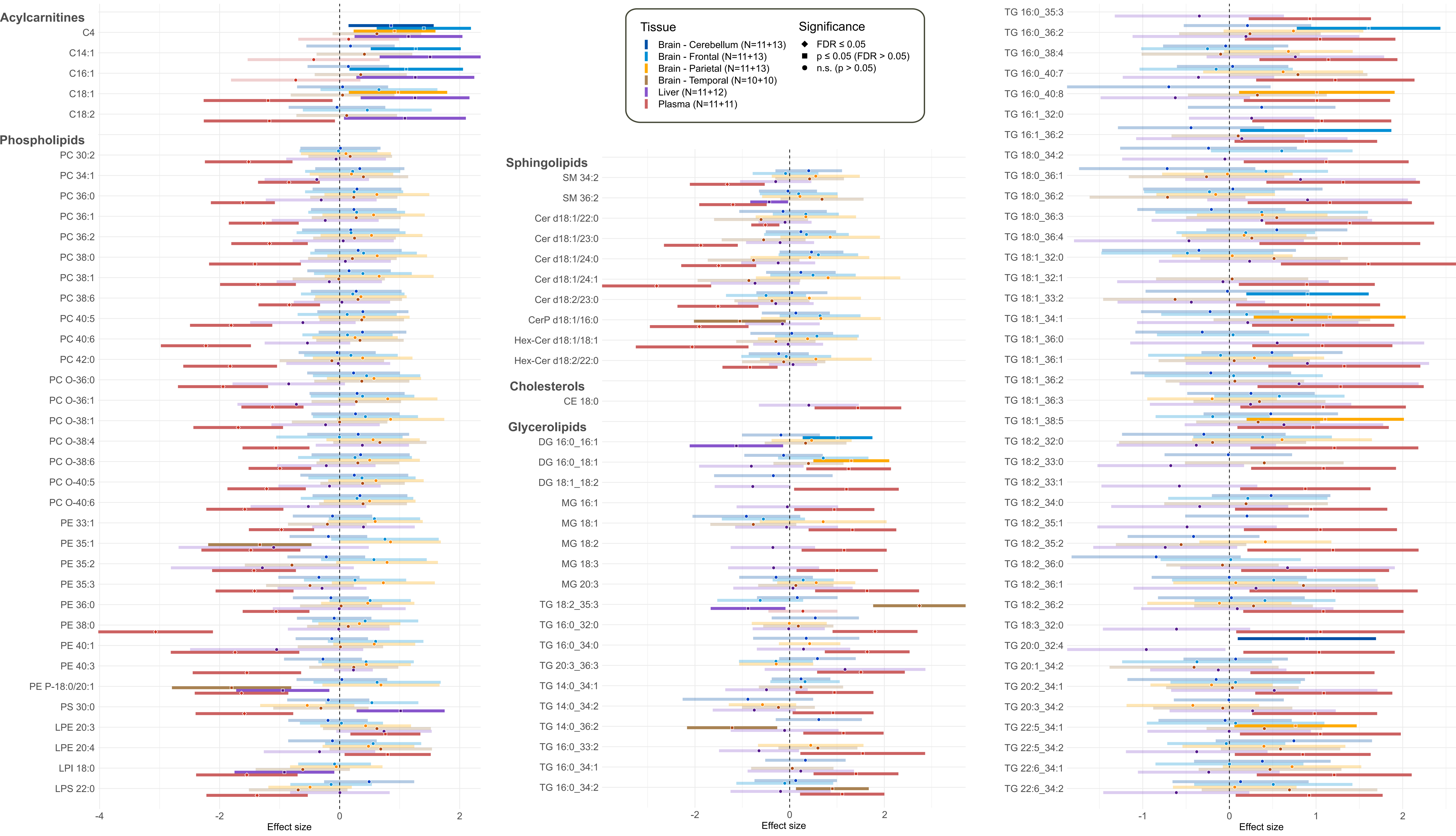
